# Projection-TAGs enable multiplex projection tracing and multi-modal profiling of projection neurons

**DOI:** 10.1101/2024.04.24.590975

**Authors:** Lite Yang, Fang Liu, Hannah Hahm, Takao Okuda, Xiaoyue Li, Yufen Zhang, Vani Kalyanaraman, Monique R. Heitmeier, Vijay K. Samineni

**Affiliations:** Department of Anesthesiology, Washington University Pain Center, Washington University School of Medicine, St. Louis, MO, United States; Neuroscience Graduate Program, Division of Biology & Biomedical Sciences, Washington University School of Medicine, St. Louis, MO, United States

## Abstract

Single-cell multiomic techniques have sparked immense interest in developing a comprehensive multi-modal map of diverse neuronal cell types and their brain-wide projections. However, investigating the complex wiring diagram, spatial organization, transcriptional, and epigenetic landscapes of brain-wide projection neurons is hampered by the lack of efficient and easily adoptable tools. Here we introduce Projection-TAGs, a retrograde AAV platform that allows multiplex tagging of projection neurons using RNA barcodes. By using Projection-TAGs, we performed multiplex projection tracing of the cortex and high-throughput single-cell profiling of the transcriptional and epigenetic landscapes of the cortical projection neurons in female mice. Projection-TAGs can be leveraged to obtain a snapshot of activity-dependent recruitment of distinct projection neurons and their molecular features in the context of a specific stimulus. Given its flexibility, usability, and compatibility, we envision that Projection-TAGs can be readily applied to build a comprehensive multi-modal map of brain neuronal cell types and their projections.

## Main

Understanding brain function requires the elucidation of the complex wiring diagram and constituent cell types across brain regions. Revolutionary work from Golgi and Cajal laid the foundation for understanding the diversity of neurons and their anatomical connections based on morphological features^2,3^. Over the past few decades, advancements have led to the further classification of neuronal cell types incorporating additional modalities, including molecular, electrophysiological, morphological, and anatomical features^4–20^. Recent breakthroughs in high-throughput single-cell techniques and spatially-resolved molecular assays have sparked immense interest in developing a comprehensive multi-modal map of diverse neuronal cell types and their brain-wide projections^21–28^. Despite the rapid integration of novel multiomic technologies in studying brain-wide connectivity, the investigation of spatially mingled neuronal projections continues to be hampered by the lack of broadly available tools to simultaneously trace multiple neuronal projections and/or profile projection neurons for multi-modal investigations.

Traditional neuroanatomical tracing methods, performed often with tracers or with viral vectors use fluorescence as the projection identifier, have been invaluable in mapping distinct neuronal projections^29–37^. However, these methods are limited by the spectral properties of fluorophores that can be detected in a single experiment, thus limiting the number of neuronal projections that can be examined simultaneously. While the recent advance in fluorescence microscopy enables simultaneous imaging of up to 10 fluorophores^38^, common microscopy equipment in neuroscience labs can reliably detect only 3-4 fluorophores, limiting the number of projections to be examined simultaneously^27,39,40^. Furthermore, such approaches are not directly suited for high-throughput sequencing-based assays such as single-cell RNA sequencing (scRNA-seq) and single-cell assay for transposase-accessible chromatin with sequencing (scATAC-seq), as the detection of exogenous transcripts is usually low by short-read sequencing. Alternatively, one can still build a multi-modal atlas for projection neurons by tracing one or two projections at a time using fluorophores and incorporating this tracing paradigm with fluorescently-activated cell sorting (FACS) for single-cell profiling^41,42^. Methods, such as retro-seq and epi-retro-seq, characterized the molecular features of projection neurons^22,26,28,43–45^, but they are often inefficient, costly, and not suitable for studying the complex wiring diagrams of projections to multiple targets.

RNA barcode-based tracing tools provide a promising avenue for high-throughput neuroanatomical studies, as diverse RNA barcodes can be parallelly detected via sequencing or imaging, thus offering a very powerful and scalable strategy to trace neuronal projections. Employing an anterograde tracing scheme, MAPSeq and BARseq utilize a sindbis viral library to encode a diverse collection of short random RNA barcodes, which are strategically designed to label individual cells and engineered to be anterogradely transported to axonal terminals for projection tracing^46–50^. Despite the ability to quantitatively measure the projection strength to any target regions of investigation, the requirement of highly customized instruments and pipelines by those methods has restricted their adoption beyond several expert labs. The cellular toxicity associated with the sindbis virus also poses a challenge to integrating their use with chronic experimental paradigms^47,51^. More recently, employing a retrograde tracing scheme using adeno-associated viruses (AAVs), both Projection-seq and MERGE-seq used RNA barcodes as the projection identifier for multiplex projection tracing^52,53^. The low cellular toxicity of AAVs allowed them to correlate gene expression to projections using scRNA-seq, but their compatibility for additional modalities, such as epigenetic profiling, spatial analysis, and activity-dependent circuit mapping is lacking or not sufficiently demonstrated.

Here we introduce Projection-TAGs, a retrograde AAV platform that allows multiplex tracing of projection neurons using RNA barcodes, which we call Projection-TAGs. Similar to Projection-seq and MERGE-seq, the key component of Projection-TAGs is a set of engineered retrograde AAVs each expressing a unique RNA barcode TAG, which acts as the projection identifier (Extended Data Fig. 1a). With this scheme, neurons projecting to a target region are uniquely labeled by a retrograde AAV-mediated TAG and multiplex projection tracing can be achieved by injecting a unique Projection-TAG AAV into each of the target regions predefined for investigation. We developed a toolbox for Projection-TAG detection using various commercial assays, enabling multiplex spatial neuroanatomical studies, high-throughput multi-modal profiling of projection neurons, and investigation of activity-dependent populations in response to a stimulus of interest. In this study, we utilized Projection-TAGs to study the complex wiring diagram, spatial organization, transcriptional, and epigenetic landscapes of intracortical-, subcortical-, and corticospinal-projecting neurons in the cortex and identify activity-dependent recruitment of distinct cortical projections in a mouse model of visceral pain. Projection-TAGs are designed for incorporation into existing experimental pipelines with minimal effort and easy application in studying the nervous system.

## Results

### Design of Projection-TAGs

To democratize and facilitate the use of Projection-TAGs in neuroscience labs without any specialized equipment, we incorporated the following features into the Projection-TAG plasmid design: First, we employed a chicken beta-actin (CAG) promoter to enable ubiquitous Projection-TAG expression in various tissue and cell types (Fig. 1a). Second, we reasoned that the inclusion of a fluorescent marker could enable flexibility in downstream processes and confirming viral expression. We screened several fusion fluorescent proteins for the ability to label both intact cells and nuclei in suspension (Extended Data Fig. 1b, c), and included in our AAV plasmids a fusion of Sad1 And UNC84 Domain Containing 1 (Sun1) to Green Fluorescent Protein (GFP), allowing enrichment of target cell/nucleus populations using FACS (Extended Data Fig. 1e). Sun1-GFP labels the nuclear membrane without interfering with the chromosome structure and has been employed in numerous epigenetic studies (Extended Data Fig. 1d)^44,54^. To further enhance Projection-TAG functionality, we developed a protocol for photobleaching Sun1-GFP fluorescence (Extended Data Fig. 2f, h) and created a version of Projection-TAGs expressing a red fluorophore oScarlet (Fig. 1a, Methods). Third, to enable projection tracing using various commercial assays, we cataloged 50 unique 100-nt RNA barcodes^55^ (Extended Data Fig. 2a, Supplementary Table 1, Methods). By insertion into the 3’ untranslated region (UTR) of the *Sun1-GFP* transcript, we generated 12 plasmids each expressing a unique Projection-TAG (TAGs 1-12, Fig. 1a). This design aimed to optimize its detection by 3’ end RNA-seq assays and enable the development of fluorescent *in situ* hybridization (FISH) probes for spatial analysis.

**Figure 1.**
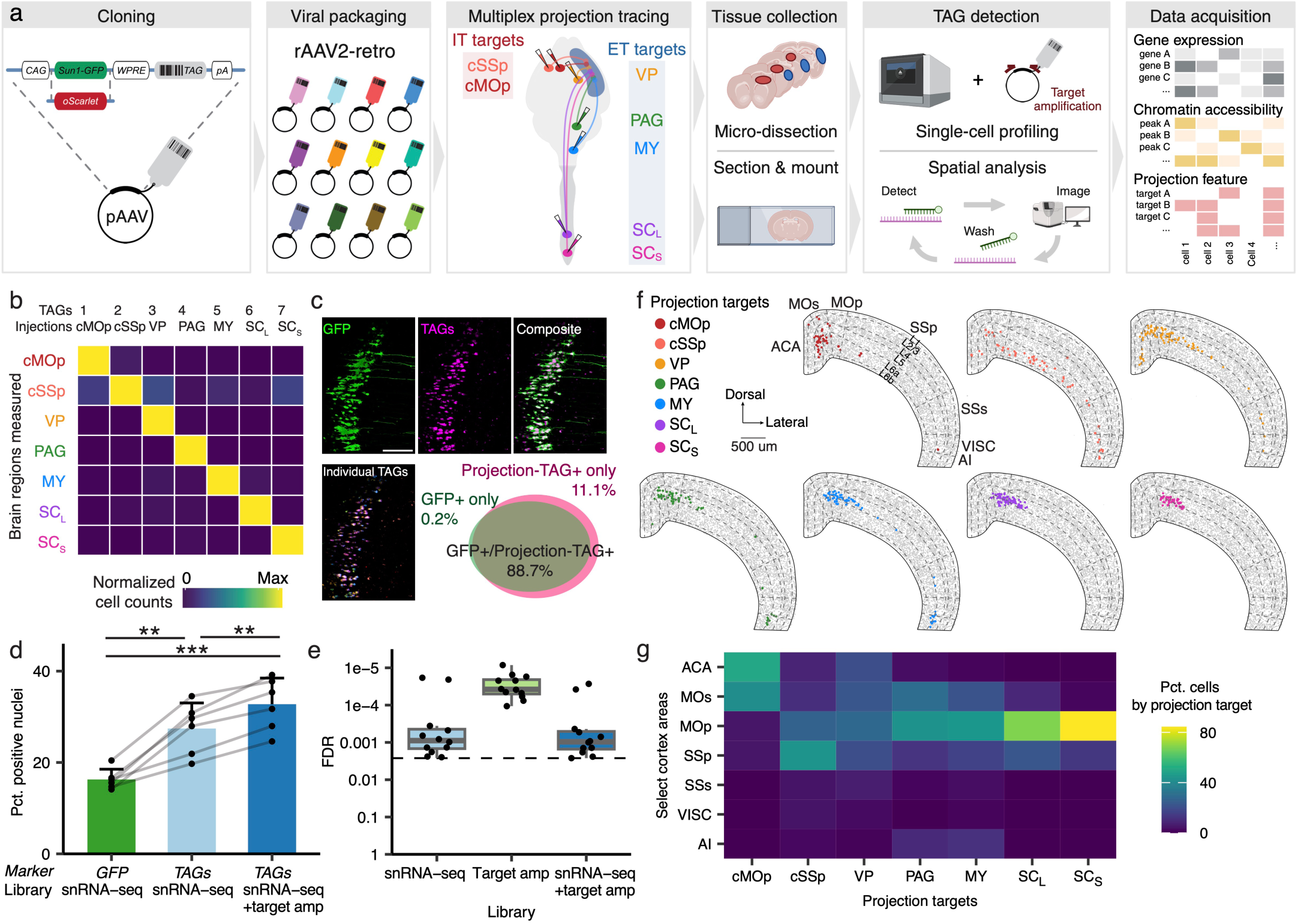
Multiplex projection tracing using Projection-TAGs. **a**, Experimental and analytical workflow of tagging neuronal projections of the MOp and SSp using Projection-TAGs. **b**, Heatmap showing the enrichment of cells labeled with individual Projection-TAGs by multiplexed FISH in brain regions of injections. Cell counts are normalized by the max values in each brain region. Individual TAGs and their injection sites are shown as on the column. **c**, Representative MOp images showing cells labeled with the fluorescent Sun1-GFP signal and fluorescent FISH signals from Projection-TAGs. Quantification shows the overlap between cells that are GFP+ and/or Projection-TAG+. FISH probes uniquely targeting Projection-TAGs (BCs 1-7) were visualized in separate imaging channels, shown as individual TAGs, and the signals were aggregated pos-hoc, shown as TAGs. Scale bar = 100 µm. **d**, Bar plot showing the percentage of nuclei expressing select marker genes (>0 UMIs for snRNA-seq, > cutoffs for target amplification, see Methods) in individual snRNA-seq libraries prepared by FACS and target amplification. Bar and error bar indicate the average and standard deviation across libraries. (*p* = 5.3e-06, *F*(2,10) = 51.77, one-way ANOVA; \*\**p* = 0.006 between Projection-TAGs and *GFP* in snRNA-seq, \*\**p* = 0.003 for Projection-TAGs between snRNA-seq only and snRNA-seq with target amplification, \*\*\**p* = 0.001 between *GFP* in snRNA-seq and Projection-TAGs in snRNA-seq with target amplification, post-hoc t-tests with Bonferroni correction). **e**, Box plot showing the false detection rate of Projection-TAGs due to ambient RNA contamination in snRNA-seq libraries. Dotted line shows the highest FDR due to ambient RNA contamination at 0.0026 among all libraries. **f**, Spatial map of Projection-TAG+ cells in select cortex areas from a brain section (−0.65 mm relative to bregma). Cells are downsampled to 200 per Projection-TAG and colored by the injection site of the corresponding TAG. Grey dots are cells segmented based on DAPI signal. **g**, Heatmap showing the distribution of projection neurons in select cortex areas from Fig. 1f. ACA: anterior cingulate area, MOs: secondary motor area, MOp: primary motor area, SSp: primary somatosensory area, SSs: supplemental somatosensory area, VISC: visceral area, AI: agranular insular area.

As a proof of principle to validate Projection-TAG expression and detection, we first performed RNA-seq and multiplexed FISH on human embryonic kidney (HEK) cells transfected with individual Projection-TAG plasmids (Extended Data Fig. 2b, d). We faithfully detected Projection-TAGs with high specificity, as the detection of Projection-TAGs expressed in each HEK sample is at least 422.4 times (RNA-seq) and 6.9 times (multiplexed FISH) greater than that of Projection-TAGs not expressed or the background (Extended Data Fig. 2c, e). To use Projection-TAGs for multiplex tracing of neuronal projections in vivo, we packaged these validated plasmids into recombinant AAV serotype 2 retro (rAAV2-retro)^30^ and generated a set of 12 Projection-TAG AAVs each expressing a unique Projection-TAG (Fig. 1a).

To evaluate the ability to use Projection-TAGs in high-throughput multi-modal profiling of projection neurons, we applied it to performing multiplex projection tracing in the adult mouse primary motor (MOp) and primary somatosensory cortex (SSp). Neurons in the MOp and SSp are anatomically organized by cortical layers and exhibit distinct layer-specific connectivity with other brain regions and have been extensively known to coordinate a wide range of innate and learned behaviors^43,56–58^. Tremendous progress has been made to characterize the diverse MOp and SSp cell types using single-cell multiomic approaches and create a comprehensive molecular taxonomy of projection neurons and their brain-wide projections^27,45,50,59–61^, allowing us to validate the efficacy of Projection-TAGs by comparing the data generated in this study with the ground truth data. To test Projection-TAGs in vivo, we identified seven downstream projection targets of MOp and SSp, including two intratelencephalic (IT) targets (contralateral MOp [cMOp] and contralateral SSp [cSSp]) and five extratelencephalic (ET) targets, which can be further classified into three subcortical targets (ipsilateral ventral posterior nucleus of the thalamus [VP], ipsilateral periaqueductal grey [PAG], ipsilateral medulla [MY]) and two corticospinal targets (lumbar spinal cord [SC_L_], and sacral spinal cord [SC_S_], Fig. 1a, Extended Data Fig. 5a).

### Multiplex projection tracing using Projection-TAG rAAV2-retro

While previous studies using rAAV2-retro have shown that two weeks are typically sufficient for fluorescently labeling cortical projections^21,22,43,44,62^, a multiplex tracing experiment would benefit from understanding the temporal kinetics and stability of cargo gene expression across different cortical projections, which remains largely elusive. We thus investigated the Projection-TAG expression in two cortical projections with notably long axons that may represent the upper limit of waiting time: MY, one of the longest cortical projections in the brain, and SC_S_, one of the longest cortical projections in the central nervous system. We injected a Projection-TAG AAV into either MY or SC_S_ and quantified the Projection-TAG expression overtime in the cortex using qPCR (Extended Data Fig. 3a). Projection-TAG expression is detectable as soon as one week after injection and increased at week two, consistent with previous reports. However, the expression in mice receiving MY injection increased rapidly and reached its peak at week three, whereas the expression in mice receiving SC_S_ injection increased more steadily and peaked at around week five. In both cases, expression plateaued until week ten, the endpoint of our study. While the initial peak timing may vary across projections, our results suggest that the stable Projection-TAG expression after peaking provides a flexible time window for synchronizing peak expression in different projections and coupling Projection-TAGs with other experimental paradigms.

To make sure Projection-TAGs can be unbiasedly used for multiplex projection tracing, we next investigated if significant viral competition exists among Projection-TAG AAVs that may reduce the tracing efficiency. Prior experiments showed that, in the medial part of the posterior parietal association area of the cortex (PTLp), more than 50% of neurons projecting to PAG also project to VP. We hypothesized that the competition between Projection-TAG AAVs would reduce tracing efficiency in mice receiving both VP and PAG injections compared to mice receiving only one of the injections. However, neither VP-nor PAG-TAG+ cell counts are significantly different from mice receiving either injection than mice receiving both injections, regardless of whether the injections were made simultaneously or separately (Extended Data Fig. 3d-f). Additionally, VP- and PAG-TAG UMI counts are not significantly different in most snRNA-seq libraries from nuclei expressing one of the TAGs than nuclei co-expressing both TAGs (Extended Data Fig. 3g,h). These results suggest that the Projection-TAG expression is largely unaffected by viral competition and thus can be safely used for multiplex tracing.

Moreover, we tested if Projection-TAG AAVs alter the gene expression and induce immune response in infected cells. In snRNA-seq, Projection-TAG+ nuclei (nuclei with >0 UMI for any Projection-TAGs) co-cluster well with Projection-TAG-nuclei on the UMAP, indicating little gene expression alteration due to AAV infection (Extended Data Fig. 3b). Gene ontology analysis revealed no significant immune response to viral infection in Projection-TAG+ nuclei compared to Projection-TAG-nuclei (Extended Data Fig. 3c). Therefore, rAAV2-retro stably express Projection-TAGs with minimal immune response.

### Detection of Projection-TAGs with high specificity and efficiency

Multiplex projection tracing relies on the demultiplexing of individual Projection-TAGs with various detection assays. In this section, we examined the specificity and efficiency of Projection-TAG detection in multiplexed FISH and snRNA-seq and highlighted some considerations that may potentially affect the interpretation of Projection-TAG experimental results.

To examine the specificity of Projection-TAG detection in vivo, we performed multiplexed FISH on brain sections from mice receiving Projection-TAG AAV injections into the projection targets of the cortex (Fig 1a, Extended Data Fig. 5a). We observed a high correspondence in Projection-TAG detection at the injection sites with 19.5 – 393 times greater cells labeled with Projection-TAGs injected into a given region than those injected into other regions (Fig. 1b, Extended Data Fig. 5b, c). We next compared the efficiency of Projection-TAG labeling to fluorescent labeling by examining MOp cells retrogradely labeled with Projection-TAGs in multiplexed FISH and those labeled with the Sun1-GFP fluorescence, expressed by the same Projection-TAG plasmids. We observed a high degree of overlap between Projection-TAG+ cells and Sun1-GFP+ cells (Fig. 1c, Extended Data Fig. 3i, j), despite Sun1-GFP and Projection-TAGs exhibiting distinct subcellular compartmentalization (Fig. 1c). Therefore, Projection-TAG labeling demonstrated high specificity and efficiency in multiplexed FISH, and that GFP fluorescence can serve as a surrogate to estimate Projection-TAG+ cells in vivo.

As different 3’ scRNA-seq technologies use varying strategies to capture RNA transcripts and construct sequencing libraries, which may affect the 3’ gene feature detection in sequencing, we next compared the Projection-TAG detection by splitting one nuclear resuspension sample into two scRNA-seq assays (Supplementary Table 2). The 10X Genomics assay yielded higher Projection-TAG detection and identified proportionally more Projection-TAG+ nuclei compared to that from Parse Biosciences, despite lower UMI recovery (Extended Data Fig. 4a). Therefore, we used the 10X Genomics assay for the sequencing experiments described in this study. Besides sequencing assays, sequencing setups also affect Projection-TAG detection. We targeted at least 100k reads/nucleus or 80% saturation rate for library sequencing, which recovered 85.2% of the Projection-TAG UMIs and 90.5% of Projection-TAG+ nuclei compared to sequencing at 500k reads/nucleus (Extended Data Fig. 4b). Sequencing read 2 length exceeding 75-nt had little effect on Projection-TAG detection (Extended Data Fig. 4c). To further improve detection and reduce overall sequencing cost, we devised a PCR-based protocol that target amplifies Projection-TAG UMIs from the cDNA library (Methods). Target amplification increased the discovery of Projection-TAG+ nuclei by 1.2 ± 0.1 fold and detection of positive nuclei for individual Projection-TAGs by 1.4 ± 0.6 fold per library (Extended Data Fig. 4d). Notably, target amplification recovered proportionally more Projection-TAG+ nuclei when the library is shallowly sequenced (Extended Data Fig. 4e), while maintaining high Projection-TAG detection.

To examine the efficiency of Projection-TAG detection, we performed snRNA-seq on FACS sorted nuclei, in which >95% of the nuclei are GFP+, from mice receiving Projection-TAG AAV injections into the projection targets of the cortex (Fig. 1a). We identified significantly more Projection-TAG+ nuclei (32.8 ± 5.7% with target amplification, 27.4 ± 5.6% in snRNA-seq library alone) than *Sun1-GFP+* nuclei (Fig. 1d). To investigate the specificity of Projection-TAG detection, we assessed the Projection-TAG mismatching in snRNA-seq. Projection-TAGs used in experiments are highly expressed (75.9 ± 6.6 percentile by expression ranks among all genes) while Projection-TAGs not used yielded zero counts in both snRNA-seq library and target amplification. We also assessed the false positive rate of Projection-TAG detection due to ambient RNA contamination and clustering and annotation errors. Projection-TAGs were detected in only 0.1 ± 0.08% of the empty droplets per library (Fig. 1e) and 0.32% of the non-neuronal cells. Target amplification marginally increased the average Projection-TAG detection in empty droplets and non-neuronal cells to 0.104% and 0.34%, respectively. These data suggest that in snRNA-seq, Projection-TAG detection is highly specific and more efficient compared to the detection of GFP expressed by the same plasmid. To control for false discovery due to technical artifacts, we applied the FDR at 0.34% in the downstream Projection-TAG analyses (Methods).

However, one should be mindful of several limitations when interpreting the Projection-TAG results. First, while the Projection-TAG detection in multiplexed FISH assay is highly efficient (only 0.2% GFP+ cells do not express Projection-TAGs), the false negative rate of Projection-TAG detection is nontrivial in snRNA-seq (67.2% ± 5.7% nuclei from snRNA-seq libraries prepared by FACS do not express Projection-TAGs, discussion). Consequently, a lack of Projection-TAG expression in a snRNA-seq cell should not be simply interpreted as lack of projection. In addition, technical and biological variables may introduce bias in Projection-TAG tracing efficiency. For example, the distribution of cells labeled for each projection differs across animals (Extended Data Fig. 4f), likely due to variations of stereotaxic injections, and the Projection-TAG expression is different across projection pathways (Extended Data Fig. 4g). However, the exact TAGs used to trace each projection pathway did not significantly affect their expression in most cases (Extended Data Fig. 4h), suggesting Projection-TAGs can be used interchangeably in tracing experiments.

### Spatial organization of projection neurons across the cortex

As it has been well established that neurons projecting to IT and ET targets exhibit distinct spatial distribution across the cortex^26,63^, we next sought to validate the spatial distribution of Projection-TAG labeled neurons following AAV injection into the projection targets mentioned above. We quantified the distribution of neurons projecting to each target in several neocortical areas (Extended Data Fig. 6c). We observed that neurons projecting to different targets are enriched in distinct areas across the cortex (Fig. 1f, g). For example, cMOp-projecting neurons are most enriched in the ACA, MOs, and the medial part of MOp, whereas cSSp-projecting neurons are dominantly found in MOp and SSp. While VP-, PAG-, and MY-projecting neurons are distributed across multiple cortex areas, SC_L_- and SC_S_-projecting neurons are highly enriched around MOp and SSp. The spatial distribution of projection neurons is largely consistent with previous reports^23,64–66^. It has been reported that thalamus- and MY-projecting neurons in the anterior lateral motor cortex exhibit distinct spatial distribution in layer 5 (L5)^26,67^. Similarly, we observed that VP-projecting neurons, while also found in L6, were more enriched in L5a than L5b (*p* = 0.005), whereas MY-projecting neurons were more abundant in L5b than L5a (*p* = 0.002) in the anterior part of the MOp (Extended Data Fig. 6d). Interestingly, their distinct sub-layer distribution appeared to attenuate towards the posterior MOp, accompanied by an increased proportion of neurons co-projecting to both targets. While numerous cortex areas contain neurons projecting to the seven targets examined in this study, our analysis highlights the MOp and the SSp as particularly rich in projection neurons among all cortex areas analyzed and are highly enriched with neurons projecting to each of the seven targets (Extended Data Fig. 6e).

### A multi-modal single-cell atlas of mouse cortex

We next assessed whether we could profile simultaneously the gene expression, chromatin accessibility, and projection feature of the same cells using Projection-TAGs in multiomic analysis of snRNA-seq and single-nucleus ATAC-seq (snATAC-seq, Extended Data Fig. 7a). We first performed snRNA-seq on four MOp samples and six SSp samples from mice receiving injections of Projection-TAG AAVs into seven downstream projection targets mentioned above, generating 14 libraries containing a total of 69,657 nuclei with an average of 3,621 genes per nucleus (Extended Data Fig. 7b, Supplementary Table 2). After quality control and removal of low-quality nuclei and doublets, 61,387 nuclei were retained in the snRNA-seq dataset. Notably, nuclei from individual libraries co-clustered together, indicating a low batch effect across libraries and largely shared gene expression profiles between nuclei from MOp and SSp (Extended Data Fig. 7c, d). Clustering analysis revealed 35 distinct transcriptional clusters, which were further classified into three major classes (Glutamatergic neurons, GABAergic neurons, and non-neuronal cells, Fig. 2a, Methods). We assigned each cluster to a known cell type, based on the expression of canonical marker genes (Fig. 2d,e, Methods). Previous studies revealed that different glutamatergic neuronal types, characterized with different anatomical properties such as layer distributions and projection targets, exhibited distinct gene expression profiles^27,43,60^. Following this naming convention, we assigned glutamatergic celltypes to 18 glutamatergic neuronal clusters based on their layer enrichment and projection patterns previously described: IT neurons from layers 2/3 (L2/3 IT), 4 (L4 IT), 5 (L5 IT), and 6 (L6 IT), pyramidal tract neurons from layer 5 (L5 PT), near-projecting neurons from layers 5/6 (L5/6 NP), corticothalamic-projecting neurons from layer 6 (L6 CT), and neurons from layer 6b (L6b). We defined three GABAergic neuronal celltypes based on developmental origins: medial ganglionic eminence (MGE) neurons and caudal ganglionic eminence (CGE) neurons. We also identified five non-neuronal celltypes: vascular cells, microglial cells (Micro.), astrocytes (Astro.), oligodendrocytes (Oligo.), and oligodendrocyte progenitor cells (OPC). We assigned subtypes to each cluster based on the marker gene distinctly expressed in that cluster compared to other clusters of the same celltype (Extended Data Fig. 7e, f). While we observed some variability of celltype distribution across libraries, likely due to different nuclear preparation approaches (Extended Data Fig. 7d), the gene expression profiles of individual subtypes are largely consistent with previous reports^15,26,61^ (Extended Data Fig. 7g, Supplementary Table 3).

**Figure 2.**
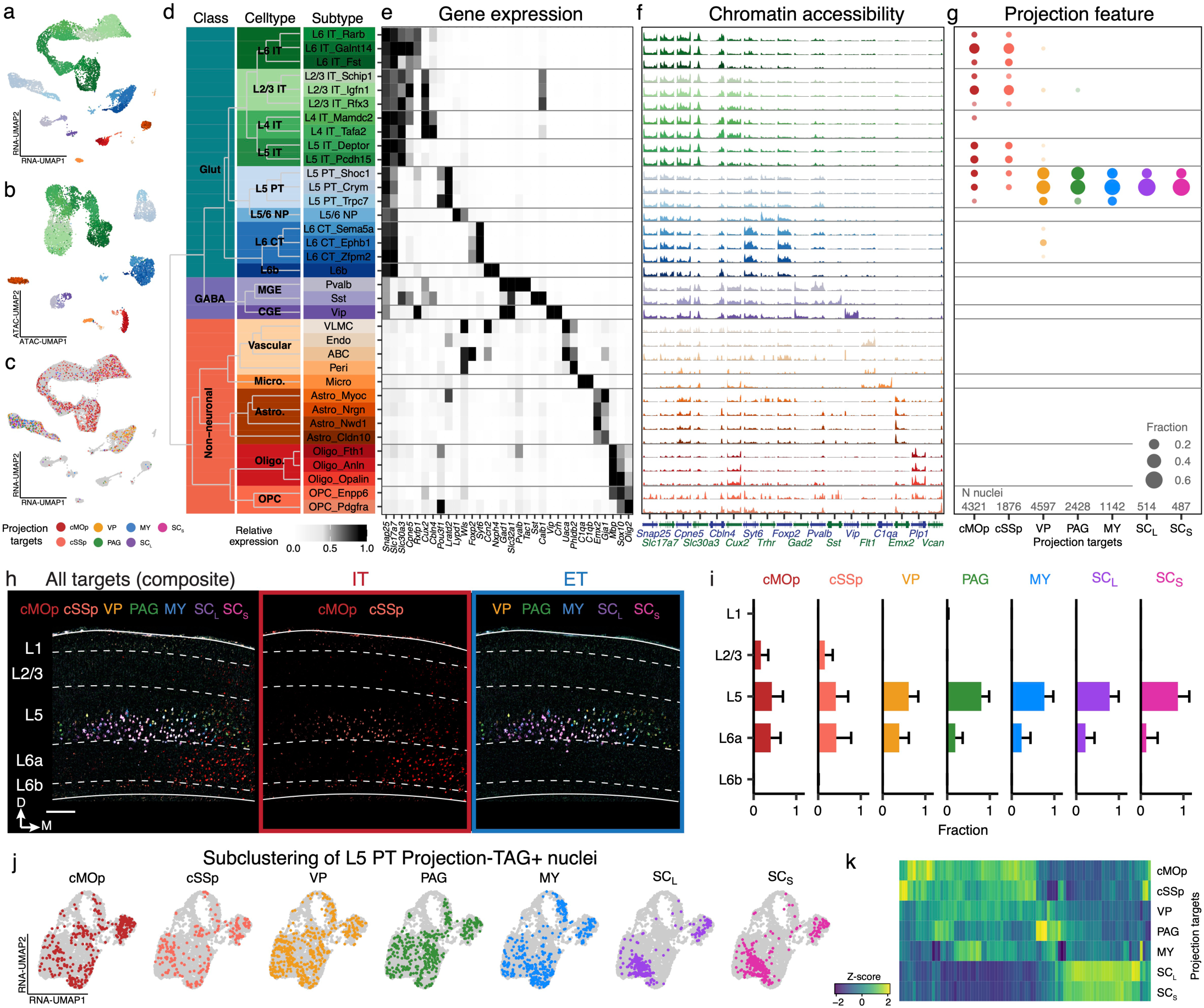
A multi-modal single-cell atlas of mouse cortex. **a**, Uniform manifold approximation and projection (UMAP)^1^ visualization of snRNA-seq data showing 10,000 downsampled nuclei, colored by transcriptional subtypes. **b**, UMAP visualization of snATAC-seq data showing 10,000 downsampled nuclei, colored by transcriptional subtypes. **c**, UMAP visualization of snRNA-seq nuclei, colored by their projection targets. 10,000 downsampled Projection-TAG-nuclei are colored grey and displayed as the background. 500 nuclei (or all nuclei if total nuclei count of one projection is less than 500) expressing only the corresponding Projection-TAG are downsampled for each projection target and displayed. **d**, Hierarchical clustering of 35 snRNA-seq clusters based on the average expression of top 100 marker genes (by FDR) per cluster, and their annotations of class, celltype, and subtype. **e**, Heatmap showing the average expression of select marker genes in individual snRNA-seq clusters. **f**, Coverage plot of average accessibility of canonical celltype-specific marker genes in snATAC-seq nuclei, grouped by transcriptional clusters. The chromatin accessibility is displayed as the average frequency of sequenced DNA fragments per nucleus for each cluster, grouped by 50 bins per displayed genomic region. **g**, Dot plot showing fraction of snRNA-seq nuclei, positive for Projection-TAG of each projection, from each transcriptional cluster. **h**, A representative FISH image showing the layer distribution of projection neurons in MOp from a brain section (−0.91 mm relative to bregma). Colors denote the projection targets based on the expression of Projection-TAGs. Scale bar = 150 µm. Left: composite image of all FISH channels, middle and right: images of a subset of Projection-TAGs injected into IT targets and ET targets, respectively. **i**, FISH quantification showing the fraction of MOp cells retrogradely labeled by each Projection-TAG in each layer. Bar and error bar indicate the average and standard deviation across six slices from two mice. **j**, RNA-UMAP showing subclustering of L5 PT neurons labeled with each Projection-TAG. **k**, Heatmap showing the z-score of average expression of genes (column) differentially expressed in L5-PT neurons positive for each Projection-TAG (row). Data shown on the heatmap can be found in Supplementary Table 4.

To enable simultaneous investigation of the gene expression and chromatin accessibility profiles in the same cells, we performed combinatorial snATAC-seq on eight snRNA-seq libraries reported above, generating a dataset of 40,188 high-quality nuclei with an average sequencing depth of 25,412 transposase-sensitive fragments per nucleus (Extended Data Fig. 8a)^68^. These snATAC-seq fragments captured the open chromatin regions of the genome as they followed the expected nucleosomal size distribution (Extended Data Fig. 8b). Clustering analysis of the snATAC-seq data revealed 28 distinct clusters. The chromatin accessibility and gene expression profiles of the sequenced nuclei are highly correlated, with 87.7 ± 18.0% nuclei in each snATAC-seq cluster assigned to the same transcriptional clusters as those made when independently analyzed for snRNA-seq data (Fig. 2b, Extended Data Fig. 7c). In addition, nuclei in individual snATAC-seq clusters display distinct chromatin accessibility around the genomic loci of canonical marker genes for the corresponding transcriptional subtypes (Fig. 2f and Supplementary Table 3). Consequently, we grouped snATAC-seq nuclei by transcriptionally defined celltypes and subtypes in subsequent analysis.

Detection and demultiplexing of Projection-TAGs by snRNA-seq enabled us to investigate the projection feature of individual sequenced cells, thus allowing multi-modal profiling of projection neurons at single-cell resolution (Fig. 2c). We determined whether a snRNA-seq nucleus project to each of the seven targets based on its expression of the corresponding TAG (Methods). To correlate the transcriptional identity with projection targets, we first examined the subtype composition of nuclei that are positive for each TAG (Fig 2f, extended Data Fig. 8d). In this analysis, a nucleus positive for the corresponding Projection-TAG will be included for the analysis of a projection target, regardless of the expression of other Projection-TAGs in the same nucleus (Methods). Hierarchical clustering based on the subtype composition for each project target divided the seven targets into three clusters (Extended Data Fig. 8e). The first cluster contains two IT targets, cMOp and cSSp. They both consist of transcriptionally-defined L2/3 IT, L4 IT, L5 IT, L6 IT, and L5 PT neurons and their subtype composition are not significantly different from each other (23.2% and 27% L2/3 IT, 4.9% and 4% L4 IT, 20.6% and 18.4% L5 IT, 29.9% and 35.8% L6 IT, 19% and 12.5% L5 PT, MOp and SSp, respectively). The second cluster contains three ET/subcortical targets: VP, PAG, and MY. While they are all enriched of L5 PT neurons (78.1% VP, 85.4% PAG, and 86.6% MY), MY-projecting neurons have proportionally fewer nuclei from L5 PT_Shoc1 cluster (20.7%, compared to 28.3% VP and 33.7% PAG). In addition, 10.4% of the VP-projecting neurons are from transcriptionally defined L6 CT subtypes. The third cluster contains two ET/corticospinal targets with SC_L_ and SC_S_ projecting neurons are enriched of L5 PT subtypes but are underrepresented in L5 PT_Trpc7 subtype (3.9% each) compared to other ET projections (subcortical: VP, PAG, and MY). Spatial analysis of multiplexed FISH further confirmed their layer distribution in MOp (Fig. 2h,i). The cell types of projection neurons are consistent with the explicit correspondence between neurons projecting to IT and ET targets previously described^26,63^.

As transcriptionally defined L5 PT neurons are enriched of nuclei positive for Projection-TAGs of all seven projection targets, we next asked whether L5 PT neurons projecting to different targets are transcriptionally distinct. Subclustering of 4,168 L5 PT Projection-TAG+ nuclei present in both snRNA-seq and snATAC-seq data revealed distinct transcriptional and epigenetic profiles of L5 PT neurons projecting to each target (Fig 2j, k, Supplementary Table 4). *Hpgd* and *Slco2a1* have been reported as marker genes for thalamus- and MY-projecting neurons, respectively^26^. Interestingly, we observed an expression gradient of those genes across L5 PT neurons projecting to different targets (Extended Data Fig. 8f), which appears to correlate to the distance of the projections, raising the possibility that the projection targets of neurons may be dictated/maintained by a shared, fine-tuned transcriptional program.

### Projection-TAGs revealed axonal collaterals and complex wiring diagram

Brain regions are interconnected with complex wiring diagrams. If a brain region projects to N downstream targets, the possible projection pattern of a single neuron will have up to 2^N^ combinations. As Projection-TAGs enable multiplex projection tracing in single animals, it opens new possibilities for studying neuronal collaterals to multiple targets. A neuron with its axonal collaterals terminated in multiple targets may be labeled by multiple Projection-TAGs. Indeed, 30.6% of Projection-TAG+ snRNA-seq nuclei express multiple Projection-TAGs (Fig. 3a). To investigate the overall pattern of axonal collaterals, we first calculated the overlap of neurons projecting to any two targets (Fig. 3b). Two IT targets (cMOp and cSSp) exhibited highly significant overlap (Extended Data Fig. 9a) with each other with 16.8% MOp-TAG+ nuclei or 36.8% SSp-TAG+ nuclei are co-labeled by both IT-TAGs. ET targets also showed highly significant overlap with each other, with the most notable overlapping to VP. 56.3 – 66.7% nuclei labeled with other ET-TAGs (PAG, MY, SC_L_, and SC_S_) are also co-labeled with VP-TAG. Additionally, PAG and MY also highly significantly overlap, as well as SC_L_ and SC_S_. To elucidate the complex projection feature of single neurons, we next investigated any possible combinations of projection to the seven targets. Out of all 128 (2^7) possible projection patterns, we identified 16 neuronal populations with distinct projection patterns that passed the FDR cutoff: seven populations with single projections (positive for only one Projection-TAG) (Fig. 3c) and nine populations with multiple projections to two or three of the targets (positive for the Projection-TAGs of the corresponding projection targets and negative for all other Projection-TAGs) (Fig. 3d). The axonal collaterals were further confirmed by multiplexed FISH (Extended Data Fig 9b). These observations are highly correlated with the anatomical hierarchy of axonal projections of the cortex previously demonstrated by anterograde bulk tracing techniques (Extended Data Fig. 9c, d). Moreover, the hierarchical organization of projections is also reflected in the gene expression and chromatin accessibility profiles of neurons projecting to individual targets (Extended Data Fig. 9e, f). This complementary analysis provides a multi-modal and high-resolution view of the spatial and molecular intricacies of neuronal projections in the mouse brain.

**Figure 3.**
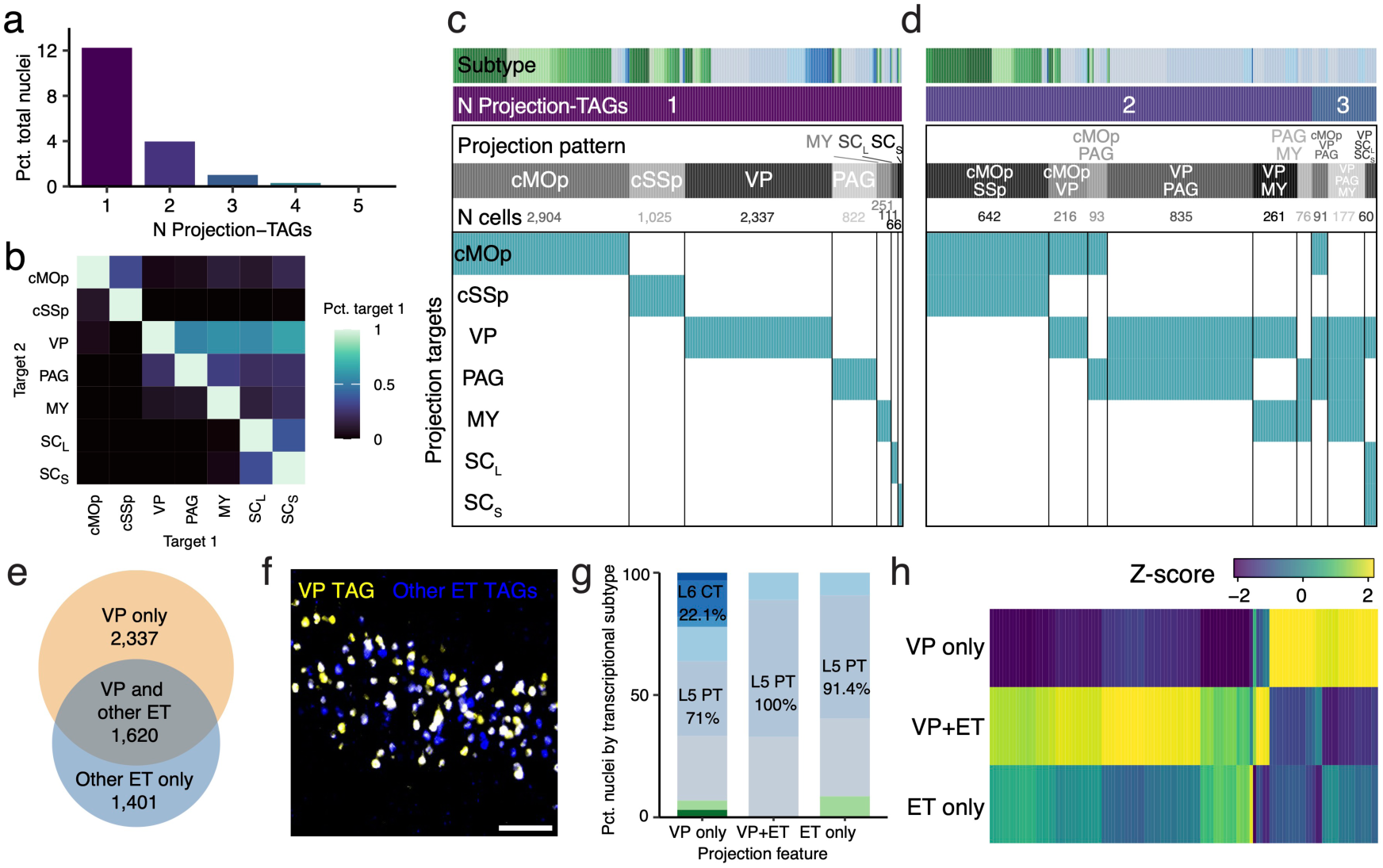
Projection-TAGs revealed axonal collaterals and complex wiring diagram. **a**, Bar plot showing the percentage of snRNA-seq nuclei, grouped by the number of unique Projection-TAGs detected in each nucleus. **b**, Heatmap demonstrating the pairwise overlapping of snRNA-seq nuclei positive for any two Projection-TAGs, shown as the percentage of nuclei positive for target 1 TAG (column) that are also positive for target 2 TAG (row). **c** and **d**, Heatmaps showing the Projection-TAG (row) detected in individual snRNA-seq nuclei (column) expressing only one Projection-TAG (c) and multiple Projection-TAGs (d). Nuclei are ordered based on the projection patterns, and only groups that passed the FDR cutoff (at least 60 nuclei) were shown on the heatmaps. **e**, Venn diagram showing the overlap between VP-TAG-expressing and other ET-TAG-expressing snRNA-seq nuclei. Other ET-TAGs include Projection-TAGs injected into PAG, MY, SC_L_, and SC_S_. **f**, Representative FISH image showing MOp cells labeled with VP- and other ET-TAGs. Scale bar = 100 µm. **g**, Distribution of transcriptional subtypes in each projection group. Transcriptional subtypes with < 60 nuclei from each projection feature group were excluded and celltypes that make up at least 10% nuclei in the projection group are annotated on the plot. **h**, Heatmap showing the z-score of average expression of 118 genes (row) differentially expressed in transcriptional L5 PT nuclei in each projection group (column). Data shown on the heatmap can be found in Supplementary Table 5.

Among 5,358 snRNA-seq nuclei positive for only ET-TAGs, 43.6% and 26.1% express only VP-TAG and only other ET-TAGs (PAG, MY, SC_L_, and SC_S_), respectively, and 30.3% express both VP- and other ET-TAGs (Fig. 3e,f). Transcriptionally defined CT neurons restrict their projection to only VP, while transcriptionally defined PT neurons broadly project to various ET targets (Fig. 3g). Our observations of axonal collaterals corroborate with the intricate single-neuron projection patterns reported in the MouseLight dataset (Extended Data Fig. 10 a,b)^69^ and further demonstrate the complexity of single-neuron projections, which has been overlooked and warrants further investigation. To uncover transcriptional programs that fine-tune cortical projection patterns, we identified 1,971 differentially expressed (DE) genes and 2,737 differentially accessible (DA) peaks in neurons with single projections compared to those with multiple projections (Fig. 3h, Supplementary Table 5).

### Characterization of genomic cis-regulatory elements and regulated genes using Projection-TAGs

While the gene expression program in the diverse MOp and SSp cell types has been largely elucidated by scRNA-seq studies, the regulatory networks that govern the distinct gene expression pattern in individual cell types and projections are under investigated^22,44,60,72^. The genomic regulatory elements (GREs), mostly identified within the non-coding region of the genome and act in a cell-type-specific and tissue-specific manner, fine tune the expression level of the regulated genes^73^. Among different types of GREs, enhancers initiate the recruitment of transcription complexes and drive the transcription of regulated genes while the silencers prevent the expression of regulated genes^74,75^. Identification of cell-type-specific GREs, specifically enhancers, has gained increased attention in the neuroscience field as they can be used as a valuable tool to mediate transgene (e.g eYFP, ChR2, DREADDS, etc) expression in the target cell populations for basic science research and potentially the development of novel therapeutics^76–81^. Though recent progress has been made in characterizing the landscape of GREs for the projection neurons, the current experimental workflow is tedious and pain-staking as individual projection neurons are labeled via injection of retro Cre in floxed nuclear reporter mice, and regions of interest are pooled for analysis of potential GREs^22,44^. We tested whether Projection-TAGs could potentially be used to perform high throughput analysis of identifying projection-specific GREs.

To this end, we integrated the snRNA-seq and snATAC-seq data from MOp and SSp to produce a unified, multi-modal cell census. We first identified 166,540 peaks when aggregated across all snATAC-seq libraries. We believe many of those peaks are likely to contain functionally relevant GREs, as 85.6% of the peaks are located at the distal regions in the genome (Fig. 4a), and 43.9% of them are differentially accessible in individual transcriptional celltypes and/or neurons projecting to different targets (Fig. 4b). To identify putative GREs and their correspondence to genes they regulate, we first identified 67,541 D peaks (Log2FC > 1, FDR < 0.05) in transcriptional celltypes and 30,843 DA peaks in neurons projecting to each target, we then calculated the average expression of genes from snRNA-seq data and the average accessibility of DA peaks from snATAC-seq data in each celltypes or projections. We next calculated the Pearson’s correlation between the accessibility of DA peaks with the expression of genes for any peak-gene pairs that are within a 5M bp window of the same chromosome (Fig. 4c). A peak-gene pair with a strong positive correlation (Pearson’s r > 0.75) is identified as a putative enhancer (pu.Enhancer) and its putative regulated gene, whereas a pair with strong negative correlation (Pearson’s r < -0.75) is categorized as a putative silencer (pu.Silencer) and its regulated gene. We identified 18,088 pu.Enhancer-gene pairs and 2,739 pu.Silencer-gene pairs that are celltype-specific (Fig. 4d,e, Supplementary Table 6), and 3,545 pu.Enhancer-gene pairs and 4,200 pu.Silencer-gene pairs that are projection-specific (Fig. 4f,g, Supplementary Table 6). Several pieces of evidence support the authenticity of identified putative GREs. First, they exhibited a greater overlap with the DNase hypersensitive sites (96.4%) detected in the adult mouse brain compared to randomly selected genomic regions (42.7%) of similar sizes and GC contents^82^. Second, these putative GREs show significant alignment with GREs previously annotated by the ENCODE consortium, particularly those identified in the mouse brain as opposed to other organs (Extended Data Fig. 11a)^83^. Furthermore, our analysis reveals four experimentally validated functional enhancers that coincide with peaks identified in our snATAC-seq data^77^. Notably, two of these validated enhancers overlap with the celltype-specific pu.Enhancers identified in our analysis, and the predicted transcriptional celltypes aligned with the cell types demonstrating the highest enhancer activity as verified experimentally (Extended Data Fig. 11b).

**Figure 4.**
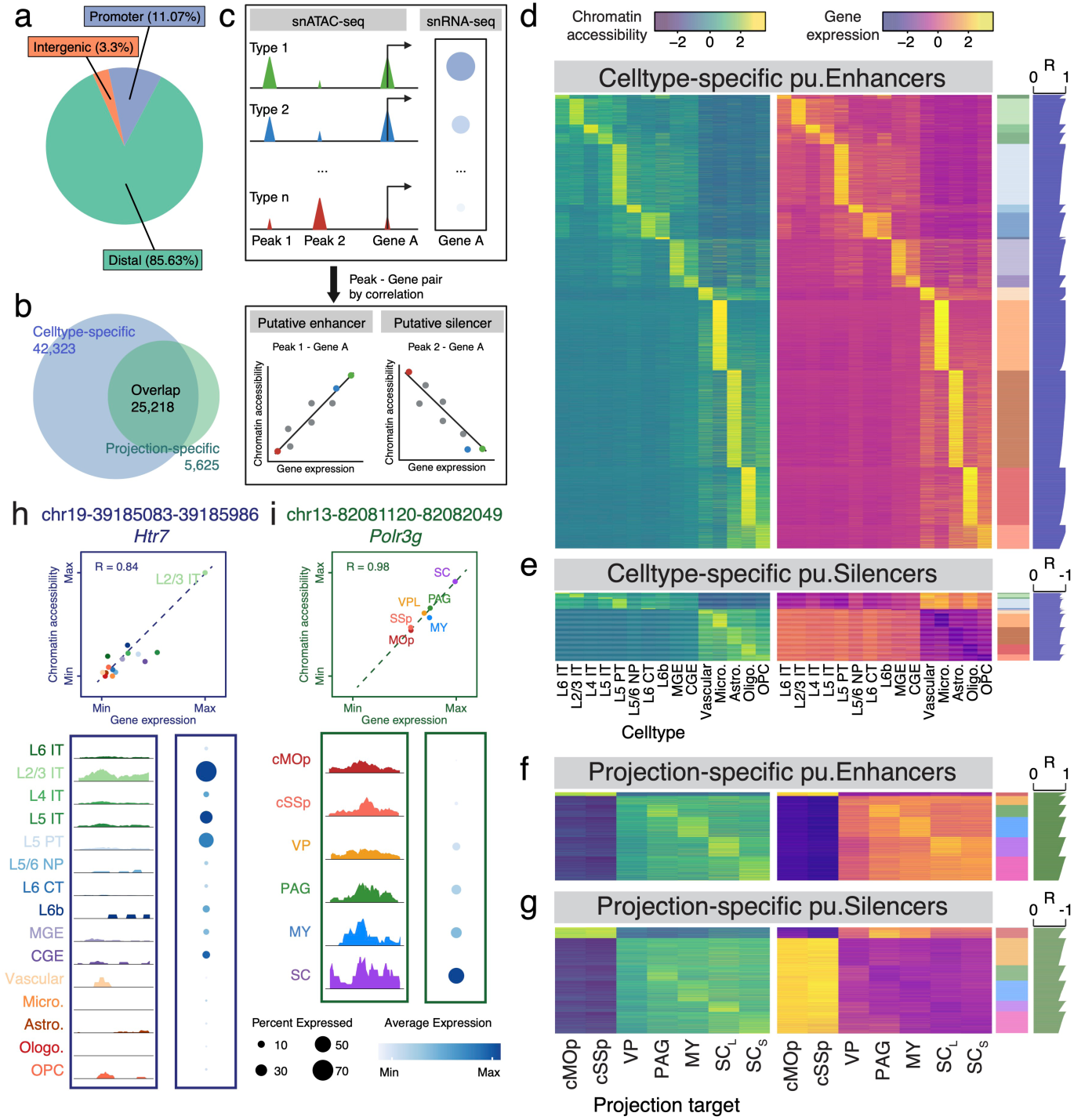
Genomic regulatory networks of cortical cell types using Projection-TAGs. **a**, Pie chart categorizing snATAC-seq peaks based on their genomic locations. Peaks are mapped to one of the three categories: promoter regions (−1,000 bp to +100 bp of transcription start site [TSS]), distal regions (<200 kb upstream or downstream of TSS or within gene body, excluding promoter), and intergenic regions (>200kb upstream or downstream of TSS, excluding gene body). **b**, Overlapping of celltype-specific peaks and projection-specific peaks in snATAC-seq data. **c**, Analytical workflow of identifying putative enhancers and silencers and their regulated genes. **d** and **e**, Heatmaps showing 18,088 pairs of celltype-specific pu.Enhancers and regulated genes (d) and 2,739 pairs of celltype-specific pu.Silencers and regulated genes (e). Left heatmaps show the average accessibility of pu.Enhancer peaks (row) in individual celltypes (column) in snATAC-seq data and right heatmaps show the average expression of regulated genes (row) in individual celltypes (column) in snRNA-seq data. On the right side next to the heatmaps, the color bar indicates the celltypes in which the peak and gene have highest accessibility and expression, and the bar plot shows the Pearson correlation coefficient (R) between the accessibility of the peak and the expression of the gene in individual celltypes. Data shown on the plots can be found in Supplementary Table 6. **f** and **g**, Heatmaps showing 3,545 pairs of projection-specific puEnhancers and regulated genes (f) and 4,200 pairs of projection-specific pu.Silencers and regulated genes (g). Left heatmaps show the average accessibility of pu.Enhancer peaks (row) in nuclei positive for each Projection-TAG (column) in snATAC-seq data and right heatmaps show the average expression of regulated genes (row) in nuclei positive for each Projection-TAG (column) in snRNA-seq data. On the right side next to the heatmaps, the color bar indicates the projection targets in which the peak and gene have highest accessibility and expression, and the bar plot shows the Pearson correlation coefficient (R) between the accessibility of the peak and the expression of the gene in nuclei positive for each Projection-TAG. Data shown on the plots can be found in Supplementary Table 6. **h**, Top: Scatter plot showing the correlation between the average accessibility of peak chr19-39,185,083-39,185,986 in snATAC-seq and the average expression of gene *Htr7* in snRNA-seq in individual celltypes (chromatin accessibility and gene expression are normalized to their max values). The dotted line represents the line of best fit. Bottom left: chromatin accessibility at the genomic locus of chr19-39,185,083-39,185,986, displayed as the average fraction of transposase-sensitive fragments per nucleus in each celltype (grouped by 50 bins per displayed genomic region). Accessibility at each locus (y axis) is scaled to the max value across all cell types. Bottom right: expression of *Htr7* in each celltype. **i**, Top: Scatter plot showing the correlation between the average accessibility of peak chr13-82,081,120-82,082,049 and the average expression of gene *Polr3g* in nuclei positive for each Projection-TAG (chromatin accessibility and gene expression are normalized to their max values). Bottom left: chromatin accessibility at the genomic locus of chr13-82,081,120-82,082,049, displayed as the average fraction of transposase-sensitive fragments per nucleus in nuclei positive ach Projection-TAG (grouped by 50 bins per displayed genomic region). Bottom right: expression of *Polr3g* in nuclei positive for each Projection-TAG. Colors indicate the injection site of the corresponding TAG. Nuclei positive for SC_L_- and SC_S_-TAGs are combined for visualization due to low cell number.

Among identified putative GREs, the peak 39,185,083 - 39,185,986 on chromosome 19 is ∼3.2Mbp upstream of the TSS of *Htr7* gene (Fig. 4h). This peak, differentially accessible in L2/3 IT neurons, is positively correlated (Pearson’s r = 0.84) to *Htr7*, which is differentially expressed in the same celltype, suggesting that this peak may contain a pu.Enhancer that might drive the expression of *Htr7* specifically in L2/3 IT neurons. The accessibility of peak 82,081,120 - 82,082,049 on chromosome 13 is positively correlated to the expression of *Polr3g* in neurons of different projections (Fig. 4i), Both the peak and gene are highly accessed/expressed in corticospinal neurons, suggesting this peak may contain a pu.Enhancer that might drive the expression of *Polr3g* specifically in the corticospinal projection neurons. Additionally, peaks chr19-10,784,771-10,785,580 and chr18-39,362,615-39,363,500 may contain celltype-specific and projection-specific pu.Silencers, respectively. While accessible in the genome, they may reduce the expression of *Fth1* in the IT neurons and *Kctd16* in ET-projecting neurons, respectively (Extended Data Fig. 11c, d). Thus, Projection-TAGs offer a powerful, high-throughput platform to perform systemic multiomic analyses to gain insight into the gene expression and chromatin accessibility profiles of diverse projection neurons.

### Projection-TAGs enable the detection of projection neurons tuned to a behavioral stimulus

Projection-TAGs allow us to elucidate the spatial, gene expression, and chromatin accessibility profiles of diverse projection neurons at single-cell resolution. As Projection-TAG AAVs mediate stable BC expression with minimal immune response, we sought to leverage it to obtain a snapshot of activity-dependent recruitment of distinct projection neurons and their molecular features in the context of a specific stimulus. The MOp and the SSp circuitry have been extensively implicated in the modulation of distinct pain modalities^84–89^, we wanted to gain insight into the gene expression changes in the cell types and their projections neurons in response to visceral pain stimulus. To study the cell populations acutely activated by visceral pain, we first labeled the seven projection targets of the MOp and SSp as described earlier (Fig. 1a). On the day of experiment, we induced acute inflammatory visceral pain with cyclophosphamide (CYP) or inject saline as the control, followed by combinatorial snRNA-seq and snATAC-seq of the MOp and SSp 30 minutes after the stimulus, at which we observed significant spontaneous behaviors (Extended Data Fig. 12a).

In both snRNA-seq and snATAC-seq data, nuclei from CYP- and saline-treated mice co-cluster together, suggesting visceral pain did not significantly alter the gene expression or chromatin accessibility of MOp and SSp cell types at the acute time point (Extended Data Fig. 12b). To pinpoint the cell populations that are acutely activated by visceral pain, we next applied Act-seq^90–92^, an approach that links the transcriptional activity following stimulus to individual neuronal cell types and projections based on the expression of immediate early genes (IEG). To aggregate expression across IEGs, we generated an IEG score for each snRNA-seq nucleus (Methods). While the IEG scores did not significantly differ in all nuclei between treatments (Extended Data Fig. 12c), CYP significantly activated 12 transcriptional subtypes, including seven IT subtypes and two GABAergic subtypes (Extended Data Fig. 12d), suggesting their preferential recruitments following the acute visceral pain stimulus. Further analysis of the projections suggests that CYP significantly activated only IT projections (both cMOp and cSSp), while none of the ET-projecting populations exhibited significant activation (Fig. 5a). Our results pinpointed the cell types and projections that visceral pain selectively recruits in the acute phase of the pain induction state.

**Figure 5.**
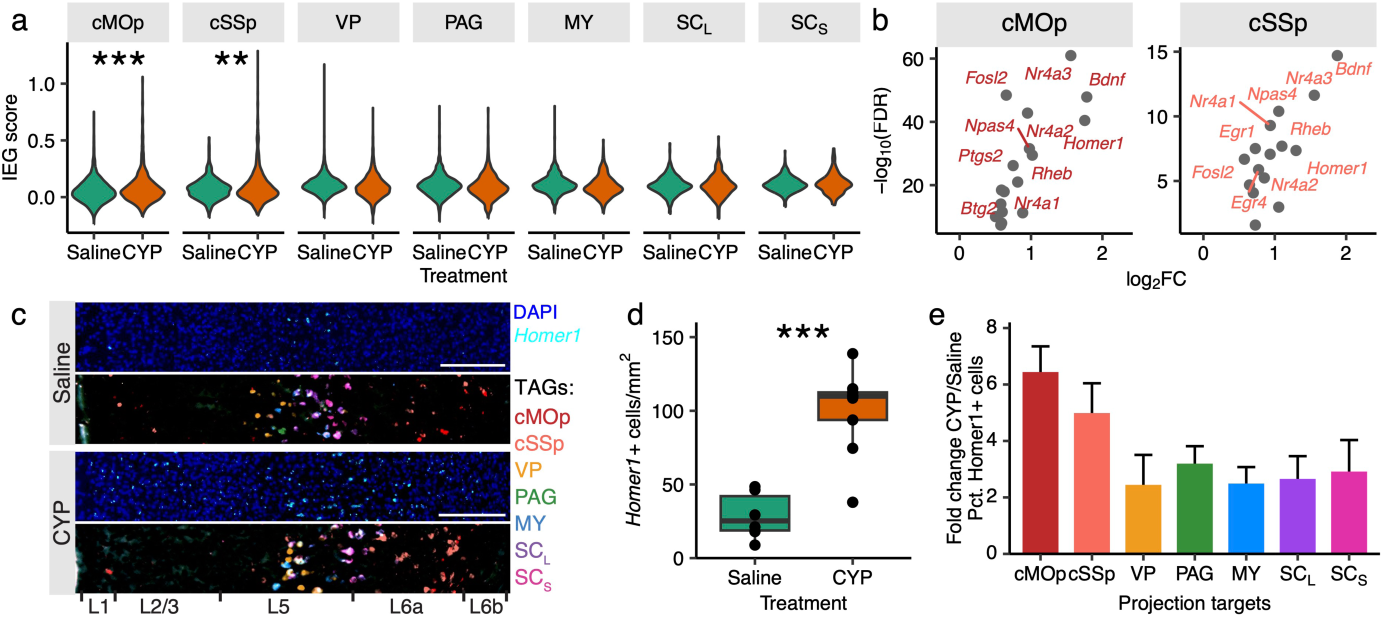
Projection-TAGs enable the detection of projection neurons acutely activated by visceral pain. **a**, Violin plots showing IEG scores in snRNA-seq nuclei for each projection between treatments (\*\*\**p* = 4.6e-14 for cMOp, \*\**p* = 0.004 for cSSp, *p* = 1 for VP, *p* = 0.99 for PAG, *p* = 0.99 for MY, *p* = 0.4 for SC_L_, *p* = 0.2 for SC_S_, one-side t-test with Bonferroni correction). **b**, Volcano plots showing significantly induced IEGs (log2FC > 0.5) in activated nuclei from CYP-treated mice compared to the same number of randomly sampled nuclei (with matched transcriptional subtypes) from saline-treated mice for the two IT projections. **c**, Representative FISH images showing the expression of *Homer1* and Projection-TAGs in MOp (from brain section approximately -0.2 mm relative to bregma) from saline- and CYP-treated mice. Scale bar: 100 µm. **d**, Quantification of *Homer1*+ cells in MOp and SSp areas from CYP- and saline-treated mice (0 to -0.92 mm relative to bregma, nine slices from three mice per group, \*\*\**p* = 1.9e-5, one-side t-test). **e**, Percentage of *Homer1*+ MOp and SSp cells in each projection. Fold change is calculated as the percentage in individual CYP slices, divided by the average percentage across saline slices. Bar and error bar indicate the mean and standard deviations across nine slices per group.

We next expanded our analysis to determine the IEG profiles of the activated IT-projecting neurons, as it has been recently appreciated that distinct stimuli exhibit varying IEG activation profiles^89,93–96^. DE analysis comparing all nuclei of the same projection between treatments failed to identify any CYP-induced IEGs. To maximize “signal-to-noise” ratio, we first identified transcriptionally “activated” nuclei based on their IEG scores (Methods), followed by DE analysis comparing activated nuclei from CYP-treated mice to inactivated nuclei from Saline-treated mice. We found that CYP induced 17 IEGs in activated neurons projecting to IT targets (Log2FC > 0.5 in either cMOp or cSSp, Fig. 5b). Among them, *Nr4a3*, *Bdnf*, *Rheb*, and *Homer1* showed higher fold change (Log2FC > 1) in both cMOp- and cSSp-projecting neurons. Interestingly, CYP did not significantly induce *Fos* expression (Fig. 5b, Extended Data Fig. 12e). *Homer1* expression is directly linked to neuronal activity and synaptic plasticity^97–100^. To validate these findings, we performed multiplexed FISH screening for *Homer1* as an activity-dependent marker for the CYP-induced pain state. We observed that CYP increased *Homer1* expression in MOp and SSp compared to saline (Fig. 5c,d). Next, to investigate if *Homer1* is selectively enriched in the IT-projecting neurons following visceral pain, we examined the colocalization of *Homer1* with each Projection-TAG. While CYP induced *Homer1* expression in all projection pathways analyzed (Extended Data Fig. 12f), IT-projection neurons exhibited a higher magnitude of *Homer1* induction, as the percentage of *Homer1*+ cells increased by 6.4 and 5-fold in cMOp and cSSp-projecting neurons, compared to on average 2.7-fold in the ET-projecting neurons (Fig. 5e). In summary, our data shows that Projection-TAGs could be readily implemented with transcriptional analysis such Act-seq to identify the molecular markers and recruitment of distinct specific brain-wide projections and cell types following the stimulus of interest.

## Discussion

Here we have developed Projection-TAGs, a retrograde AAV platform for multiplex neuroanatomical studies and high-throughput multi-modal profiling of projection neurons. Projection-TAG AAVs retrogradely label distinct projections with unique RNA barcodes with high specificity and efficiency. It can be easily adaptable to existing neuroscience workflow and is optimized for commercial assays, such as multiplexed FISH, which allows simultaneous spatial integration of multiple projections in the same animals, and single-cell profiling assays that enable multiomic profiling of projection neurons. In this study, we applied Projection-TAGs to examine the transcriptional and epigenetic landscapes of the cortex using combinatorial snRNA-seq and snATAC-seq. Projection-TAGs could facilitate the development of novel tools for projection-specific targeting/manipulation by opening new avenues for studying multiple projections in the same animals and for identifying key gene expression features and genomic regulatory elements in distinct cortical cell types and diverse projection neurons. Lastly, we demonstrated that Projection-TAGs can be incorporated with additional experimental paradigms, providing users with flexibility for studying activity-dependent recruitment of distinct cell populations and projections in the stimulus of interest.

### Available high-throughput neuroanatomy tools

RNA barcode-based high-throughput neuroanatomical tools have greatly expanded the multiplexing capacity of neuronal tracing. Available high-throughput neuroanatomical techniques can be broadly classified based on their retrograde and barcoding schemes. Anterograde tracing techniques, namely MAP-seq and its derivatives, such as BRICseq, BARseq, and BARseq2^46,49,50,101^. MAPseq utilizes a sindbis viral library to encode a diverse collection of short random RNA barcodes, which act as the cell barcode/identifier. The sindbis viral library is injected into the source region, and a unique barcode is expressed in individual neurons and anterogradely transported to the axonal terminals. Multiplexed projection tracing is achieved by assaying the RNA barcodes present in each of the target regions. MAPseq can be coupled with scRNA-seq for multi-modal profiling of projection neurons^48^ and with in-situ sequencing for spatial analysis (BARseq) and spatially-resolved transcriptional assays (BARseq2). Due to the anterograde tracing scheme, BARseq and BARseq2 can achieve high spatial resolution in both source and target regions, but the multiplexing capacity and tracing accuracy of MAPseq may be limited by tissue dissections. With the simple surgery procedure (one injection into the source region), MAPseq and derivatives are ideal for quantitatively measuring the projection strength. However, the requirement of highly customized instruments and pipelines by those methods has restricted their adoption beyond several expert labs. The high replication rate of sindbis virus results in cellular toxicity, posing a challenge to integrating their use with chronic experimental paradigms.

Retrograde tracing techniques, including Projection-seq, MERGE-seq, and Projection-TAGs^53,102^. These techniques utilize a limited number of RNA barcodes with known sequences, which act as the projection barcode/identifier. A retrograde AAV expressing a unique BC is injected into each of the target regions, which retrogradely label projection neurons in the source region. Multiplexed projection tracing is achieved by assaying the barcodes present in the source region, which can be simply read out by scRNA-seq and spatial assays. Due to the retrograde tracing scheme, Projection-TAGs and similar techniques can achieve high spatial resolution in the source region, but not necessarily the target regions. The surgery procedures are relatively complicated (one injection into each of the target regions), which may limit the multiplexing capacity, and the accuracy of targeting each target region may introduce technical variations that confound the quantitative measurement of the projection strength. Surgical targeting (one injection into each of the target regions) may introduce technical variations that confound the quantitative measurement of the projection strength and might limit the multiplexing capacity. Despite these technical challenges, barcode detection is relatively simple and flexible and can be achieved using various commercial assays. Projection-TAGs enable multiplex neuroanatomical studies and high-throughput multi-modal profiling of projection neurons. AAVs have minimal cellular toxicity, which is ideal for incorporating Projection-TAGs with additional experimental paradigms for studying activity-dependent recruitment of distinct cell populations and projections in the stimulus of interest.

### Consideration for Projection-TAG experimental design and result interpretation

Projection-TAGs allow multiplex projection tracing and multi-modal profiling of projection neurons. In retrograde viral tracing experiments, the cargo gene expression in a single cell depends on a series of biological processes such as viral attachment and internalization, transduction and trafficking to the cell body, and escape and entry into the nucleus^103^. rAAV2-retro has been widely adopted in neuroscience laboratories, Projection-TAG rAAV2-retro performance would be similar to other rAAV2-retro viruses, and prior experience using rAAV2-retro may be used to guide the planning of Projection-TAG multiplex tracing experiment. Efficient and accurate labeling of Projection-TAGs relies on successful targeting of desired brain regions, transduction efficacy of rAAV2-retro, titer (Projection-TAG AAVs with the titer of ∼2e+12vg/ml label 14.7 ± 8.4% of MOp cells, compared 3.7 ± 3.0% by those with the titer of ∼5e+11vg/ml) and volume of the viruses injected. The high correlation between Projection-TAG (RNA level) and GFP (protein level) expressed using the Projection-TAG AAVs provides a simple strategy for investigators to estimate retrograde tracing efficiency and accuracy of target labeling, which are essential for a successful Projection-TAG experiment. While Projection-TAG plasmids have the potential to be flexibly packaged into other AAV serotypes, the transduction efficiency and tissue/celltype tropisms related to AAV serotypes may lead to false negatives and should be considered while designing the experiments and interpreting the results. While Projection-TAG AAVs can be multiplexed and used interchangeably (Extended Data Fig. 3d-h, Extended Data Fig. 4h), AAV transduction efficiency and Projection-TAG expression may not be uniform across projection pathways (Extended Data Fig. 4f, g), which may introduce confounds to quantitative measurement of the projection strength.

After the tracing experiments, Projection-TAG detection and multi-modal profiling can be achieved using various commercial assays. Projection-TAG detection is specific and sensitive in single-cell sequencing and multiplexed FISH, as previously described (Fig. 1). Multiplexed FISH is superior at the Projection-TAG detection efficiency but is labor-intensive and difficult to scale up. Sun1-GFP expressed by Projection-TAG AAVs reliably labels both whole cells and nuclear suspensions, enabling enrichment of projection neurons for unbiased single-cell and single-nucleus transcriptional and epigenetic profiling in a high-throughput manner. Enzyme-free nuclear extraction works for various tissue samples (fresh, frozen, or fixed) and introduces minimal dissociation-induced transcriptional stress response, and thus ideal for activity-dependent circuit mapping. Whole-cell sequencing has the ability to sequence both cytoplasmic and nuclear transcripts, advantageous for recovering medium-to low-expressing transcripts and may potentially increase Projection-TAG detection. Sequencing involves many biochemical processes that may confound the Projection-TAG detection. Artifacts such as ambient RNA contamination, doublets, and analytical errors may lead to false positives, which can be addressed experimentally (e.g. reduce ambient RNA contamination by adding additional wash steps and incorporating FACS) and computationally (e.g. ambient RNA correction, doublet removal, and setting up appropriate FDR for the analysis). Projection-TAG detection rate is limited by the sequencing assays (e.g. efficiency of reverse transcription and cDNA capture) and sequencing setups (Extended Data Fig. 4), which may pose an upper limit for Projection-TAG detection and lead to false negatives. Consequently, lack of Projection-TAG expression in a cell should not be simply interpreted as lack of projection, and the neurons projecting to each target may be under-reported in snRNA-seq, which may result in a higher degree of underestimation of the axonal collaterals. Projection-TAG detection using sequencing assays with higher transcript capture efficiency and multiplexed FISH (or multi-color CTB tracing) may circumvent false negatives.

### Future directions of Projection-TAGs

Projection-TAGs could potentially be compatible with additional commercial platforms, such as high-throughput spatial transcriptomics platforms like Xenium and Visium, that would allow spatially-resolved investigation of diverse cell types and distinct anatomical organization of the projection neurons. Another potential application of Projection-TAGs includes connecting neuronal activity to the distinct projections by integrating the oScarlet version with in vitro and in vivo calcium imaging experiments. The cell type and projection information from imaging-based molecular assays can be integrated with the real-time neuronal activity information from the calcium imaging experiments via post-hoc imaging alignment and registration^93,104–106^.

The number of projections that can be labeled with Projection-TAG AAVs is not inherently constrained, and Projection-TAGs could be easily scalable as we have screened 50 BCs that could potentially be packaged to increase multiplexing of projection tagging. It has been well appreciated that AAV serotypes have distinct tropism and selective labeling of the brain nuclei. With the discovery of capsid selection using directed evolution^107–109^ and advances in sequencing techniques and computational tools to optimize exogenous transcript detection^110,111^, we anticipate that the improved retrograde labeling efficiency of viruses will enhance wider applications of Projection-TAGs. Given its flexibility, usability, and compatibility with commercial platforms, we envision that Projection-TAGs can be readily applied to study diverse projection types in the central and peripheral nervous system.

## Methods

### Generation of Projection-TAGs

We used 100-bp BCs because they allow us to design HCR probes to detect the spatial distribution of BC transcripts, and would improve their detection rate in RNA-seq compared to shorter BCs that are commonly used. We first retrieved 60 previously reported BCs^55^. We then filtered out the BCs that contain the sequence of restriction enzymes (BamHI, HpaI, and NotI). Next, we performed sequence alignment using blastn suite searching in the “Nucleotide Collection (nr/nt)” database and “refseq_representative_genomes” database against the genomes of human (taxid:9606), mouse (taxid:10090), rat (taxid:10116), and Primates (taxid:9443) and filtered out the ones that showed significant similarities^112^. We then calculated the Euclidean distance between any of the two BCs using DistanceMatrix() function in R package DECIPHER. We then cataloged and reported the first 50 BCs sorted based on the average Euclidean distances with other BCs.

### Cloning and viral packaging of Projection-TAG AAVs

To generate the backbone of pAAV-CAG-Sun1-GFP-WPRE-pA, we first linearized two previously reported plasmids: pAAV-CAG-H2B-GFP (Addgene, Plasmid #116869)^26^ using HindIII and pAAV-Ef1a-DIO-Sun1GFP-WPRE-pA (Addgene, Plasmid #160141)^113^ using AscI, and end filled. We then digested them with SpeI and NheI, respectively, followed by gel extraction of the fragments at around 4,700 bp (pAAV-CAG backbone) and 2,400 bp (Sun1-GFP), respectively. We next generated the plasmid pAAV-CAG-Sun1-GFP-WPRE-pA by ligating the two fragments together. To insert BCs into pAAV-CAG-Sun1-GFP-WPRE-pA, we first generated a gene block fragment for each BC with the following structure (5’ overhang-NotI-BC-SV40 polyA-HpaI-3’ overhang) and sequence (AAGGAAAAAAgcggccgc-100bp sequence of BC-ttcgagcagacatgataagatacattgatgagtttggacaaaccacaactagaatgcagtgaaaaaaatgctttatttgtgaaatttgtgatgctattgctttatttgtaaccattataagctgcaataaacaagttaacAACCGCTGCCG). We then digested both pAAV-CAG-Sun1-GFP-WPRE-pA and BC gene blocks with NotI and HpaI, and generated pAAV-CAG-Sun1-GFP-WPRE-BC-pA by ligating the fragments. We generated 12 plasmids each expressing a unique BC (1 - 12). We packaged the plasmids into AAV (BrainVTA, China) and generated a set of 12 rAAV2-retro samples each expressing a unique BC (titer range 1.1 - 3.5e+12 vg/ml). To generate a set of 12 plasmids pAAV-CAG-oScarlet-WPRE-BC-pA each expressing a unique BC, we generated BamHI-oScarlet-WPRE-NotI gene fragment and replaced Sun1-GFP-WPRE fragment in pAAV-CAG-Sun1-GFP-WPRE-BC-pA using BamHI and NotI. All restriction enzymes were purchased from New England Biolabs, and all plasmids produced in this study are deposited to Addgene: https://www.addgene.org/browse/article/28247961/.

### Testing Projection-TAGs in HEK cells

We acquired human embryo kidney cell line HEK 293T/17 (ATCC) and cultured it using DMEM (Corning), supplemented with 10% Fetal Bovine Serum (Gibco), and 1% Penicillin/Streptomycin (Sigma), on either 60 mm dishes for RNA-seq or 8-well chambered slides (Nunc™ Lab-Tek™) pre-coated with PDL-collagen for multiplexed FISH. Once the cells reached 70 – 80% confluency, we transfected the DNA of one BC plasmid into each of the HEK samples using lipofectamine 2000 (Invitrogen) according to the manual. Cells were incubated for 24 – 36 hours before analysis. For RNA-seq, around 1 million cells were used for each sample and RNA was extracted and purified using RNAqueous™ Total RNA Isolation Kit (Invitrogen) following the manual. Library preparation and sequencing were conducted by the McDonnel Institute of Genomics at Washington University School of Medicine. For FISH, cells were washed with DPBS and fixed using 4% formaldehyde for 10 minutes at room temperature. FISH was performed following the manufacturer’s manual. See the following section “Multiplexed fluorescent *in situ* hybridization with HCR” for experimental details.

### Animals

All experiments were conducted in accordance with the National Institute of Health guidelines and with approval from the Animal Care and Use Committee of Washington University School of Medicine. Mice were housed on a 12-hour light-dark cycle (6:00 am to 6:00 pm) and were allowed free access to food and water. All animals were bred onto C57BL/6J background, and no more than five animals were housed per cage. Female littermates between 8 and 10 weeks old were used for experiments.

### Stereotaxic surgeries

Mice were given a single subcutaneous injection of 0.5ml 0.9% sterile sodium chloride and Buprenorphine-SR, 1 hour prior to surgery to help rehydrate the mouse after anesthesia. Mice were anesthetized with 1.5 - 2% isoflurane in an induction chamber using isoflurane/breathing air mix. Once deeply anesthetized, mice were secured in a stereotactic frame (RWD Life Science) where surgical anesthesia was maintained using 2% isoflurane. Mice were kept on a heating pad for the duration of the procedure. Preoperative care included application of sterile eye ointment for lubrication, administration of 1 mL of subcutaneous saline, and surgery-site sterilization with iodine solution. We injected into the seven projection targets of the cortex, each with a Projection-TAG (rAAV2-retro expressing a unique TAG), in single animals (Supplementary Table 2). All injections were made using a Nanoject II auto injector (Drummond) with a glass microelectrode at a rate of 1 nl/s and the needle was held in place for 10 minutes prior to needle withdrawal. Needles are changed between each Projection-TAG injection to avoid contamination between Projection TAGs. Stereotaxic surgery injecting into SC_L_ and SC_S_ was performed at week 0. For SC_L_ injection, the injections were located 400 μm lateral to the center of the posterior artery (150–300 mm below the dura), and the virus was bilaterally injected between T12 and T13 intravertebrally with 250 µl volume each injection. For SC_S_ injection, the dorsal part of the L1 vertebra was gently excised using a high-speed micro-drill (RWD Life Science) to unveil the L6-S1 spinal cord segments. The injections were located 100 μm left-lateral to the center of the posterior artery with a 10-degree left-right tilt (550–580mm below the dura). Three distinct sites were injected with 150 nl of virus at each site. Following a 2-week convalescent interval, the second stereotaxic surgery was conducted by injecting into cMOp, cSSp, VP, PAG, and MY using the following coordinates relative to Bregma: cMOp (anterior-posterior [AP] -0.52 mm, medial-lateral [ML] +1.81 mm, dorsal-ventral [DV] -0.86 mm, left hemisphere), cSSp (AP -0.52 mm, ML +0.75 mm, DV -0.86 mm, left hemisphere), VP (AP -1.5 mm, ML -1.52 mm, DV -4 mm, right hemisphere), PAG (AP -4.5 mm, ML -0.5 mm, DV -2.83 mm, right hemisphere), and MY (AP -6.2 mm, ML -0.5 mm, DV -5.93 mm, left hemisphere). A small midline dorsal incision was performed to expose the skull. After leveling the head relative to the stereotaxic frame, the specified injection coordinates were used to mark the locations on the skull, and a small hole (approximately 0.5 mm diameter) was drilled for each. 500 nl of virus was injected into each of the targets. After each surgery, the surgery site was sutured, and mice were recovered from anesthesia on a heating pad and then returned to their home cage. Mice were given carprofen (0.05 mg/ml in water) to minimize inflammation and discomfort and monitored for three consecutive days.

To label neurons co-projecting to cSSp and VP, 500 nl of AAV2retro-hSyn-eGFP-Cre (Addgene #105540) was injected into VP and 500 nl of AAV2retro-Ef1a-DIO-H2B-tdTomato (BrainVTA) into cSSp of the same mice (n=2) using the stereotaxic coordinates described above. Retrograde labeling in the SSp and MOp was examined two weeks after injection.

### Mouse model of visceral pain

Stereotaxic surgeries were performed as described above. At week 5 post-viral injections, mice were acclimated for at least 3 days prior to the experiment. Mice were administered an intraperitoneal injection of either cyclophosphamide (Sigma-Aldrich, dissolved and diluted to 40 mg/mL in 0.9% sodium chloride) with a dose of 200 mg/kg or saline (approximately the same volume as CYP). Animals were monitored in their home cage for 30 minutes before perfusion.

### Tissue processing

In the experiments reported in this paper, we traced the seven projection targets of the MOp and SSp by performing two stereotaxic surgeries using undiluted AAVs in 8-10 weeks-old C57BL/6 female mice. Individual Projection-TAG AAVs were injected into the SC_L_ and SC_S_ at week zero, followed by a second surgery of AAV injections into the MOp, SSp, VP, PAG, and MY at week two. Animals were perfused, cortical samples were collected for either FISH or sequencing analysis at week five.

To collect biological samples for FISH studies, animals were anesthetized with ketamine cocktail, and perfused with DEPC-PBS followed by 4% DEPC-PFA. Brain and spinal cord tissues were dissected and incubated in 4% DEPC-PFA at 4 C for 6 - 8 hours. Tissues were then incubated in 30% sucrose in 1X DEPC-PBS at 4 C for 24 - 48 hours until they sank to the bottom of the tube. Tissues were then embedded in OCT and stored at -80 C° before slicing. Tissues were sliced coronally into sections with 30 µm thickness using a cryostat (Leica), and tissue slices were mounted on microscope slides. Slides were stored at -20 C° for a least one hour before moving to -80 C° for long-term storage. Slides containing the brain regions of injections were examined under a fluorescent microscope, and the fluorescent signal from Sun1-GFP was used to confirm the injection sites. As an optional step, the native fluorescence of Sun1-GFP can be photobleached from mouse brain sections by floating slices in 1×PBS with 24 mM NaOH and 4.5% H_2_O_2_ and exposing under the UV light (27 total watts, OPPSK) for 30 minutes at room temperature. Slices were immediately rinsed with DEPC-PBS twice before mounting on a microscope slide.

To collect biological samples for molecular experiments such as qPCR or sequencing, animals were anesthetized with ketamine cocktail, and perfused with ice-cold NMDG-based cutting solution (NMDG 93 mM, KCl 2.5 mM, NaH_2_PO_4_ 1.25 mM, NaHCO_3_ 30 mM, HEPES 20 mM, Glucose 25 mM, Ascorbic acid 5 mM, Tiourea 2 mM, Sodium Pyruvate 3 mM, MgSO_4_ 10 mM, CaCl_2_ 0.5 mM, N-acetylcysteine 12 mM; pH adjusted to 7.3 with 12N HCl, and bubbled with 95% O_2_ and 5% CO_2_). The spinal cord was dissected, fixed, and sectioned by following the sample preparation process mentioned above in order to confirm the injection sites of SC_L_ and SC_S_. The brain was submerged in ice-cold NMDG-based cutting solution and sliced coronally into sections with 400 µm thickness using a Compresstome (Precisionary, VF-210-0Z). MOp and SSp samples were prepared by micro-dissecting the respective regions under a microscope and collected into a nucleus-free centrifuge tube placed on dry ice. Samples were stored at -80 C° for long-term storage.

To assess the viral spread in the injection sites, the remaining brain and spinal cord sections containing the injection sites were placed on glass slides. The fluorescent signal from Sun1-GFP was examined under a fluorescent microscope. We first examined and confirmed the GFP expression in each of the injection sites for every mouse included in this study. To assess off-target labelling, we next examined if there was strong GFP signal displayed in the neighbor regions of the injection site that are away from the intended seed region, and if there is leaked labelling in the needle track. We quantified the off-target effect for each injection site by calculating the area of GFP signal displayed in the neighbor brain regions divided by the area of the intended target brain region (Extended Data Fig. 5). On average, the off-target effect was measured at 39.6 ± 16.4% (cMOp), 8.4 ± 5.6% (cSSp), 16.2 ± 8.5% (VP), 14.7 ± 10.6% (PAG), and 8.8 ± 6% (MY) across all mice used in this study. GFP signals in the spinal cord are restricted to the target spinal cord segment for all spinal cord injections across all mice used in this study.

### Multiplexed FISH with HCR

HCR (hybridization chain reaction) v3.0 probes, amplifiers, and reagents were purchased from Molecular Instruments^114^. Multiplexed FISH was performed according to the manual of HCR RNA-FISH with minor modifications. For hybridization, samples were equilibrated in hybridization buffer for 30 min at 37 C° and hybridized with probe sets (16 nM each probe) in hybridization buffer overnight at 37 C°. Samples were washed in probe wash buffer and gradually switched to 5×SSCT (5×SSC, 0.1%Tween-20) at 37 C°. HCR was carried out at room temperature. Samples were equilibrated in amplification buffer for 30 min and the amplifier hairpins (conjugated to Alexa-488, Alexa-546, Alexa-647, and/or Cyanine 7) were heated to 95 C° and snap-cooled in a dark drawer for 30 min. Hairpins were then mixed and diluted to 0.6 nM each hairpin in amplification buffer before incubated with samples overnight.

Samples were washed with 5×SSCT for a total of three times and imaged in 2×SSCT with DAPI. Imaging was carried out as described in the next section. After imaging, samples were washed with 5×SSCT, then HCR probes and hairpins were stripped by incubating with 0.25 U/ul of DNase I (Sigma Aldrich) in 1×DNase incubation buffer for 90 minutes at 37 C° (Extended Data Fig. 2g, i). Samples were washed with 5×SSCT for 5 minutes for a total of 5 times before the next round of HCR-FISH.

### Imaging

After each FISH round, slices were imaged at 10x magnification on a fluorescence microscope (Keyence, BZ-X800). Filter cubes used include DAPI (Chroma 49021, Excitation filter at 405 nm with 20 nm bandwidth, Emission filter at 460 nm with 50 nm bandwidth), GFP/AF488 (Chroma 49011, 480/40x, 535/50m), Alexa-546 (Chroma 49304, 546/10x, 572/23m), Alexa-647 (Chroma 49006, 620/60x, 700/75m), and Cyanine 7 (Chroma 49007, 710/75x, 810/80m). The DAPI signal and signals from FISH probes were imaged for each FISH round. Images from each FISH round were stitched using the Keyence BZ-X800 Analyzer. Images from multiple FISH rounds were loaded into FIJI/ImageJ and aligned using the HyperStackReg plugin by choosing DAPI channels for transformation matrix computation. Images were then downsampled by a factor of 2 and background was subtracted.

### Quantification of projection neurons using QUINT

FISH images were segmented and quantified using the QUINT workflow which allows for semi-automated quantification of cells in labeled brain regions^115^. For atlas registration, images from different channels were merged and downsampled according to the manual. An XML file was generated using Filebuilder and loaded into QuickNII for linear registration to the Allen Mouse Brain Atlas CCFv3^116,117^. After, user-guided nonlinear refinements were applied to brain sections using VisuAlign. For the segmentation of positive cells, images of individual channels were prepared, and segmentation was performed through the pixel classification and object classification pipelines in ilastik^118^. In pixel classification, models were trained to distinguish signal from background using a small subset of images and applied to the whole dataset. Individual machine-learning algorithms were trained for each channel. Probability maps of signal and background were exported as HDF5 files to be used in object classification. In object classification, models were trained to distinguish objects from non-specific labeling based on features such as size and shape using a small subset of images applied to the whole dataset.

The performance of illastik segmentation algorithms was validated against manual segmentation. Illastik accurately identified 96.8% of manually segmented cells with a 6.6% false positive rate (Extended Data Fig. 6b). There are 2.1% of the segmented objects identified as multiple cells merged into one object and 1.9% of the segmented objects identified as one cell split into multiple objects. Finally, segmentation and registration files were uploaded to the Quantifier feature in Nutil which resulted in quantification of cells per brain region according to the reference atlas^119^. To identify cells that co-labeled by multiple markers, objects positive for each marker were segmented individually and the x and y coordinates of the center of each object were calculated using illastik. If the Euclidian distance of two objects from separate channels is less than the average radius of all objects, the two objects were identified as one cell that co-expresses both marker genes. For Projection-TAG detection and specificity analysis, cSSp and cMOp were excluded from this analysis because they are known to contain neurons projecting to other injection sites and may be labeled by TAGs injected into other regions which could contaminate the final distribution of the TAG.

### Single-nuclei isolation and FACS

Nuclear extraction was performed according to the protocol described previously with minor modifications^120^. Mouse cortical tissues were transferred to a dounce homogenizer in homogenization buffer (0.25 M sucrose, 25 mM KCl, 5 mM MgCl2, 10 mM Tris-HCl, pH 8.0, 5 ug/mL actinomycin, 1% BSA, and 0.08 U/ul RNase inhibitor, 0.01% NP40) on ice. Samples were homogenized for 10 strokes with the loose pestle in a total volume of 1 mL, followed by 10 additional strokes with the tight pestle. The tissue homogenate was then passed through a 50 µm filter and diluted 1:1 with working solution (50% iodixanol, 25 mM KCl, 5 mM MgCl2, and 10 mM Tris-HCl, pH 8.0). Nuclei were layered onto an iodixanol gradient after homogenization and ultracentrifuged as described previously. After ultracentrifugation, nuclei were collected between the 30 and 40% iodixanol layers and diluted with resuspension buffer (1xPBS with 1% BSA, and 0.08 U/ul RNase inhibitor). Nuclei were centrifuged at 500 g for 10 min at 4°C and resuspended in resuspension buffer with 5 ng/ul of 7-AAD. For FACS, gates on GFP and 7AAD were set using tissues collected from mice with no stereotaxic surgeries and no 7AAD staining. GFP+/7-AAD+ events and GFP-/7-AAD+ events were sorted using 100 µm nozzle on a BD FACSARIA II and collected separately into 1.5 mL microcentrifuge tubes containing 100 µl of resuspension buffer.

### snRNA-seq and snATAC-seq

For snRNA-seq, nuclei were further processed and sequenced according to the manufacturer’s manuals of Parse-biosciences Evercode WT mini Kit V2 (Parse) or 10X Genomics Chromium Single Cell Gene Expression 3’ V3.1 Assay (10X RNA). For combinatorial snRNA-seq and snATAC-seq, nuclei were processed and sequenced according to the manufacturer’s manual of 10X Genomics Chromium Single Cell Multiome Assay (10X Multiome). Libraries were sequenced on a NovaSeq6000 with 150 cycles each for Read1 and Read2, targeting 100,000 paired reads/nucleus for snRNA-seq libraries and 50,000 paired reads/nucleus for snATAC-seq libraries. Raw sequencing data from individual libraries were processed and mapped using 10X Genomics cellranger-6.1.2 (10X RNA), 10X Genomics cellranger-arc-2.0.1 (10X Multiome), or Parse Computational Pipeline v1.1.2 (Parse). The reference genome was generated by adding the Sun1-GFP and Projection-TAGs 1-12 [Supplementary Table 1]) to GRCm38. Projection-TAG feature was extended from the 100-nt Projection-TAG sequence by 140-nt on each end, to ensure sequencing reads have at least 10-nt alignment to the Projection-TAGs . For snATAC-seq, accessible peaks were identified using cellranger-arc aggr by analyzing the combined fragment signals across all snATAC-seq libraries in the dataset.

### Target Amplification

Parallel PCR reaction (KAPA HiFi HotStart ReadyMix, Roche) was carried out to amplify Projection-TAG sequences from the cDNA libraries generated using 10X Genomics Chromium Single Cell Gene Expression 3’ V3.1 or Multiome Assay, First, forward primer (5’-ACACTCTTTCCCTACACGACGCTCTTCCGATCT-3’) and reverse primer (5’-GTGACTGGAGTTCAGACGTGTGCTCTTCCGATCTCCTCCCCGCATCGATACCG’-3’) was used separately in PCR reactions with 20 ng of cDNA library each reaction. PCR program: (1) 95 °C for 3 min, (2) 98° for 20 s, 70 °C for 15 s, 72 °C for 20 s (10 cycles), (3) 72 for 1 min. The reactions containing forward and reverse primers were pooled and a second PCR reaction was done following the PCR program above. As a result, the region containing the Projection-TAG sequence, 10X cell barcodes, and UMI sequences will be preferentially amplified from the cDNA molecules. The target amplification amplicons were labeled with sample indices (Dual Index Plate TT Set A, 10X Genomics) and using PCR reaction with program (1) 95 °C for 3 min, (2) 98° for 20 s, 65 °C for 15 s, 72 °C for 20 s (12 cycles), (3) 72 for 1 min. PCR products were purified using SPRIselect beads sequenced on a NovaSeq6000 with 150 cycles each for Read1 and Read2, targeting 10M reads per library. Raw sequencing data from individual libraries were processed and mapped to individual Projection-TAGs using 10X Genomics cellranger-6.1.2.

### Quality control, clustering, and annotation of snRNA-seq

The gene-cell count matrices from all snRNA-seq libraries were concatenated using R (V4.1.1) package Seurat (V4.4.0)^121^. To be included in the snRNA-seq analysis, nuclei were required to contain at least 500 unique genes, less than 15,000 total UMIs, and fewer than 5% of the counts deriving from mitochondrial genes. There were 69,657 nuclei that met these criteria. Raw counts were scaled to 10,000 transcripts per nucleus and log-transformed using NormalizeData() function to control the sequencing depth between nuclei. Counts were centered and scaled for each gene using ScaleData() function. Highly variable genes were identified using FindVariableFeatures() and the top 20 principal components were retrieved with RunPCA() using default parameters. For dimension reduction and visualization, Uniform Manifold Approximation and Projection (UMAP) coordinates were calculated using RunUMAP(). Nuclei clustering was performed using FindClusters() based on the variable features from the top 20 principal components, with the resolution set at 0.6, and the marker genes for each cluster was identified using FindAllMarkers() comparing nuclei in one cluster to all other nuclei. Doublet or low-quality nuclei were identified if they meet any of the following criteria: 1). Assigned to a cluster with no significantly enriched marker genes (FDR < 0.05, log2FC > 1); 2). Assigned to a cluster in which five or more mitochondrial genes were identified among top 20 marker genes (sorted by avg_log2FC); 3). Identified as doublets using R package DoubletFinder with doublet expectation rate at 5%^122^. In total 5,991 nuclei were identified as doublet or low-quality and thus excluded from the dataset. The remaining 63,666 nuclei were clustered as described above.

Transcriptional classes and cell types were assigned to each cluster based on the canonical marker genes previously reported (Fig. 2e). Specifically, for classes, glutamatergic and GABAergic neuronal clusters are annotated based on the expression of *Slc17a7* and *Slc32a1*, respectively. Non-neuronal clusters are annotated based on the lack of expression of *Slc17a7* and *Slc32a1*. For cell types, IT clusters are annotated based on the expression of *Slc30a3*, and PT, NP, CT clusters are labeled by *Lratd2*, *Lypd1*, and *Syt6*, respectively. L6b is marked by *Nxph4*. Vascular cells express *Crh* and/or *Uaca*. Microglial cells and astrocytes are marked by *C1qa* and *Emx2*, respectively.

Oligodendrocytes and OPC are labeled based on the expression of *Mbp* and *Sox10*. Subtypes were assigned to each cluster based on the marker gene uniquely expressed in that specific cluster (Extended Data Fig. 7,e,f).

### Projection feature annotation

To study the projection feature of single neurons, we assigned the projection targets to individual snRNA-seq nuclei based on the expression of Projection-TAGs. In snRNA-seq, a Projection-TAG-cell matrix was generated from the gene-cell matrix. A projection target is assigned to a single nucleus if that nucleus expresses the Projection-TAG (> 0 UMIs) injected into the target, and each nucleus is evaluated for each projection target based on the expression of the corresponding Projection-TAG. In target amplification, we set an expression cutoff for each Projection-TAG in each library. The cutoff is 1 UMI or X percentile of the corresponding Projection-TAG expression in the corresponding library (where the X equals the percentage of nuclei not expressing the corresponding Projection-TAG in the corresponding snRNA-seq library plus one), whichever comes greater. A projection target is assigned to a single nucleus if that the Projection-TAG expression is greater than the cutoff, and each nucleus is evaluated for each projection target based on the expression of the corresponding Projection-TAG. Finally, to combine results from snRNA-seq and target amplification, a projection target is assigned to a single nucleus if that target is assigned in either snRNA-seq or target amplification. Of note, a nucleus can be assigned multiple projection targets if it expressed multiple Projection-TAGs. In the analysis where individual projection targets are analyzed separately (e.g. Fig. 2), nuclei were grouped based on whether they express the corresponding Projection-TAG. For example, snRNA-seq nuclei only expressing cMOp-TAG and co-expressing cMOp-TAG and other TAGs will be grouped for cMOp analysis. In the analysis where axonal collaterals were reported (e.g. Fig. 3), nuclei were grouped based on the expression of each of the Projection-TAGs. For example, snRNA-seq nuclei only expressing cMOp-TAG and co-expressing cMOp-TAG and cSSp-TAG will be grouped separately for analysis. Projection groups with sample size no less than 60 nuclei (corresponding to 0.34% of the total snRNA-seq nuclei generated from FACS-sorted libraries, the upper limit of FDR described in Fig. 1) were reported in all analyses.

### Anchoring analysis of snRNA-seq

To validate the transcriptional cluster annotation, we directly compare our snRNA-seq data to a published scRNA-seq of mouse MOp and SSp previously described^123^. We used Seurat to anchor our snRNA-seq data to the published dataset. First, FindTransferAnchors() was used to identify anchors (conserved features) between datasets. TransferData() was run to transfer cell type labels described in the published dataset to each nucleus in the our snRNA-seq data.

### Quality control and clustering of snATAC-seq

The peak-cell counts matrix was loaded and analyzed using R package Signac (V1.7.0)^124^. To be included for snATAC-seq analysis, nuclei were required to be present in snRNA-seq data and contain at least 1,000 fragments overlapping with peaks. There were 41,553 nuclei that met these criteria. Term frequency-inverse document frequency normalization was perfomed using RunTFIDF () function and variable peaks were identified using FindTopFeatures (). Dimension reduction was performed with singular value decomposition using RunSVD() function and Uniform Manifold Approximation and Projection (UMAP) coordinates were calculated using RunUMAP(). Nuclei clustering was performed using FindClusters() based on the top 20 dimensions with the resolution of 0.8.

### Differential analyses of gene expression and chromatin accessibility

To identify marker genes and peaks that are enriched in distinct transcriptional subtypes or projections, differential expression/accessibility analysis was performed using findAllMarkers() in Seurat/Signac, comparing nuclei from one subtype or projection to all other nuclei. Genes and peaks with FDR < 0.05 were reported. To identify genes and peaks that are differentially expressed/accessible between any two subtypes or projections, differential expression/accessibility analysis was performed using findMarkers() in Seurat/Signac, comparing nuclei from one subtype or projection to nuclei from another subtype or projection. Genes and peaks with FDR < 0.05 were reported.

To identify genes that are differentially expressed in neurons projecting to only one target compared to neurons projecting to multiple targets while controlling for transcriptional cell types, differential expression analysis was performed using FindMarkers() in Seurat, comparing nuclei of the same transcriptional cell type expressing only one Projection-TAG to nuclei co-expressing the same Projection-TAG and other TAGs. Cell types that have at least 60 nuclei for each projection group were analyzed, and genes and peaks with FDR < 0.05 were reported.

To identify genes and peaks that are differentially regulated by CYP in individual subtypes or projections, differential expression/accessibility analysis was performed using findMarkers() in Seurat/Signac, comparing nuclei from CYP-treated animals to nuclei from Saline-treated animals of the same subtype or projection. Genes and peaks with avg_log2FC > 0.5 and FDR < 0.05 were reported. To identify IEGs that are activated by CYP in neurons projecting to IT targets, cMOp and cSSp, differential expression analysis was performed using findMarkers() in Seurat, comparing activated nuclei, defined by Act-seq, that are positive for the corresponding Projection-TAG from CYP-treated animals to the same number of randomly sampled nuclei (with matched subtype distribution) positive for the same Projection-TAG from Saline-treated animals. IEGs with avg_log2FC > 0.5 and FDR < 0.05 were reported.

### Identification of putative genomic regulatory elements

To identify snATAC-seq peaks that are correlated with gene expression in cis and may act as genomic regulatory elements (GREs) to regulate the expression of target genes, we modified a computational pipeline previously described^54^. we first calculated, using snATAC-seq data, the average accessibility of each peak and, using snRNA-seq data, the average expression of each gene in individual Subtypes or Nuclei positive for individual TAGs. We then computed the Pearson correlation coefficient r between the accessibility of peaks and the expression of genes across Nuclei positive for individual TAGs or Subtypes for any peak-gene pair, in which the center of the peak is located within 5 mbps of the center of the TSS of the gene on the same chromosome. To identify putative GREs that may direct celltype-specific gene expression, we identified pairs of Celltype-specific snATAC-seq peaks (avg_log2FC > 0.5, FDR < 0.05, comparing peak accessibility in nuclei of one Celltype to all others) and Celltype-specific snRNA-seq genes (avg_log2FC > 0.5, FDR < 0.05, comparing gene expression in nuclei of one T-type to all others) in the same T-type. A peak-gene pair with a strong positive correlation (Pearson’s r > 0.75) is identified as a putative enhancer (pu.Enhancer) and its putative regulated gene, whereas a pair with a strong negative correlation (Pearson’s r < -0.75) is categorized as a putative silencer (pu.Silencer) and its regulated gene. To identify putative GREs that may direct Projection-specific gene expression, we identified pairs of Projection-specific snATAC-seq peaks (avg_log2FC > 0.5, FDR < 0.05, comparing peak accessibility in nuclei of one Projection to all others) and Projection-specific snRNA-seq genes (avg_log2FC > 0.5, FDR < 0.05, comparing gene expression in nuclei of one Projection to all others) in the same Projection. Projection-specific pu.Enhancers and pu.Silencers and their regulated genes are identified as described above.

### Act-seq analysis

To identify cell populations that are activated by visceral pain, the IEG score for each nucleus in snRNA-seq data was calculated using AddModuleScore() in Seurat based on the expression of a set of 139 immediate early genes (IEG) previously described^90^. A transcriptional subtype or projection was considered transcriptionally “activated” if the IEG scores of CYP-treated nuclei in this population were significantly higher than the IEG scores of Saline-treated nuclei. A nucleus was considered transcriptionally ‘‘activated’’ if its IEG score was 2 standard deviations higher than the average IEG scores across all Saline-treated nuclei in the same projection group (nuclei expressing the corresponding TAG).

### Statistical analysis and visualization

Statistical analyses including the number of animals or cells (n) and *p* values for each experiment are noted in the figure legends. Statistics and visualization were performed using R version 4.0.1. Student’s t-tests were performed using R package stats V4.2.2. ANOVA tests were performed using R package rstatix V0.7.2, followed by post-hoc t-tests with Bonferroni correction. Hypergeometric tests were used to test the significance of the overlap between two gene sets or between nuclei expressing two TAGs using by calling phyper() function in R package stats V4.2.2. Plots were generated using R packages ggplot2 V3.4.0, gplots V3.1.3, and UpSetR V1.4.0.

## Data and code availability

Raw and processed data of snRNA-seq and snATAC-seq experiments included in this study are deposited to the NCBI Gene Expression (GEO) SRA with accession number <u>GSE277718</u>. Custom pipelines and scripts are available on GitHub: https://github.com/Samineni-Lab/Projection-TAGs.

## Acknowledgements

This work was funded by National Institute on Drug Abuse (R01DA056829) and National Institute of Diabetes and Digestive and Kidney Diseases (R01DK128475) awarded to V.K.S. We thank Flow Cytometry & Fluorescence Activated Cell Sorting Core in the Department of Pathology and Immunology at Washington University School of Medicine for help with FACS. We thank the Genome Technology Access Center at McDonnell Genome Institute at Washington University School of Medicine for help with snRNA-seq and snATAC-seq library construction and sequencing. The Center is partially supported by NCI Cancer Center Support Grant #P30 CA91842 to the Siteman Cancer Center from the National Center for Research Resources (NCRR), a component of the National Institutes of Health (NIH), and NIH Roadmap for Medical Research. We would like to thank Dr. Robert W Gereau IV, Dr. William Renthal, and Dr. Lisa Fang for their helpful discussion of the manuscript and experimental design; we would like to thank all the Samineni and Gereau lab members and WashU Pain Center colleagues for their help with manuscript preparation.

## Author contributions

V.K.S. and L.Y. conceptualized the project and designed the experiments; L.Y., V.K.S., and M.R.H. generated the plasmids; F.L. performed the anatomical tracing experiments; H.H., F.L., T.O., and Y.Z. collected the samples for the library preparation and *in situ* validation studies. L.Y. and H.H. performed multiplexed FISH and analyzed the data; L.Y. performed sequencing experiments; L.Y. and X.L. performed the sequencing analysis; V.K.S. and L.Y. wrote the manuscript with inputs and feedback from all authors.

## Competing interest

The authors declare that they have no conflict of interest.

---

**Extended Data Fig. 1 – related to Fig. 1.**
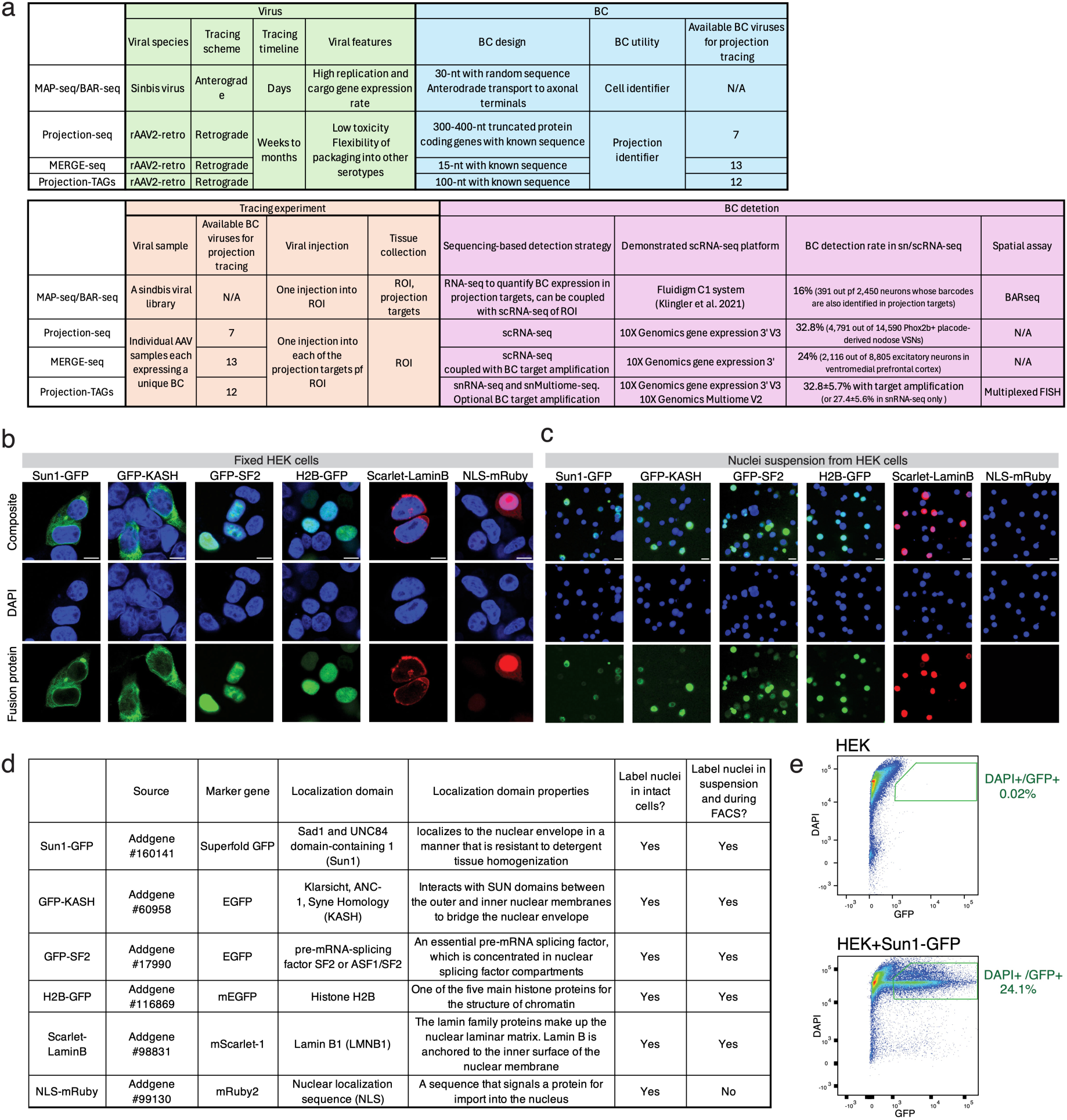
**a**. Summary of available high-throughput neuroanatomical tools. **b**, Representative fluorescent images of the localization of fusion proteins in different cellular compartments in fixed HEK cells. Scale bar: 10 µm. **c**, Representative images of the fluorescent labeling using fusion proteins of nuclear resuspension extracted from HEK cells. Scale bar: 20 µm. **d**, Summary of the fluorescent proteins and their ability of labeling of cells and nuclei of HEK cells. **e**, Scatter plot showing the gate of sorting labeled HEK nuclei (DAPI+ and GFP+) extracted from HEK cells expressing Sun1-GFP.

**Extended Data Fig. 2 – related to Fig. 1.**
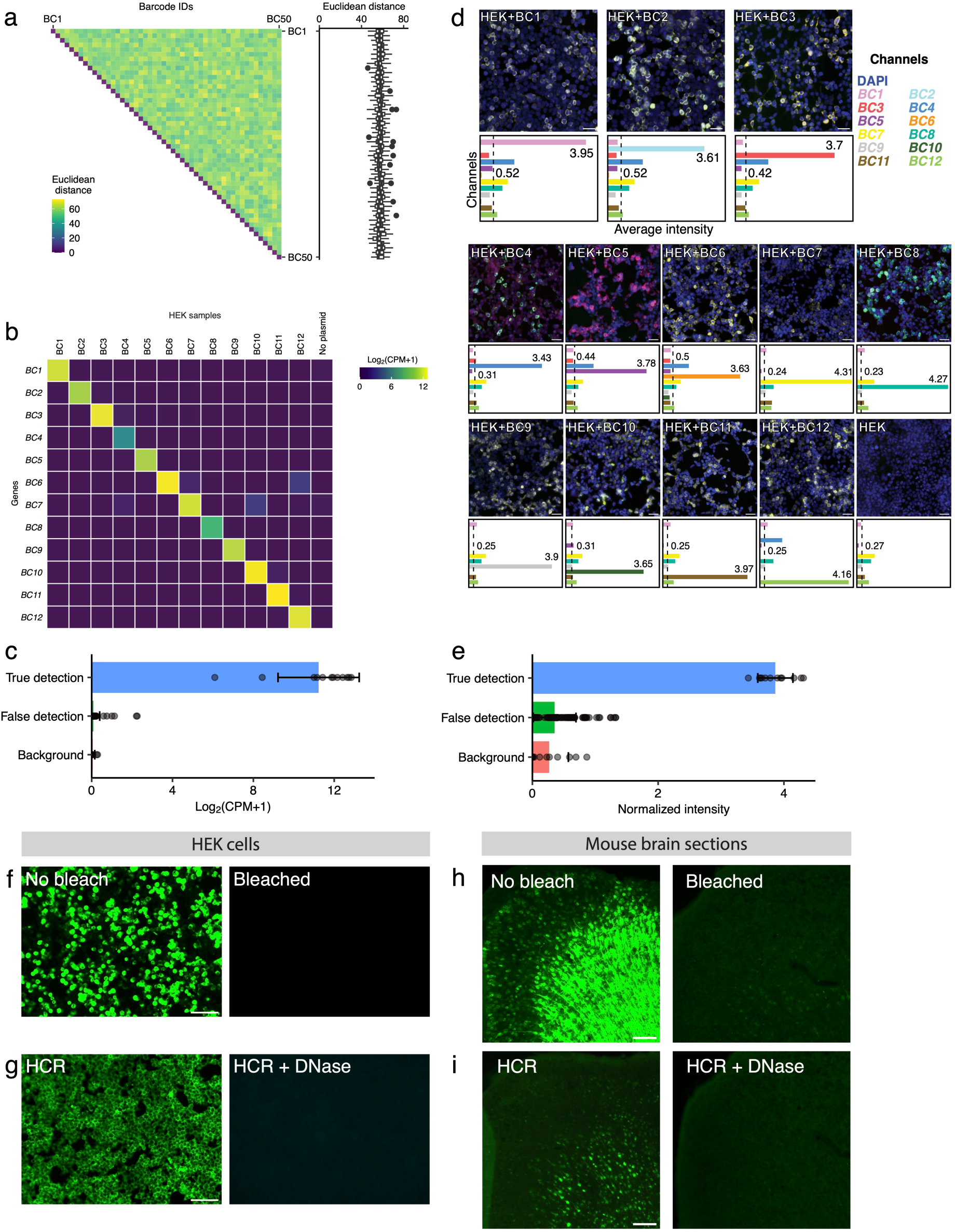
**a**, Left: pairwise Euclidean distance between any two of the 50 BCs cataloged in this study. Right: box plot showing the average Euclidean distance of individual BCs to other BCs. **b**, Counts of sequencing reads that map to individual BCs (row) in individual HEK samples (column) in RNA-seq. Each HEK sample was either transfected with a plasmid expressing a unique Project-TAG BC or no transfection. **c**, Bar plot showing the average detection rate of Projection-TAGs in individual samples, error bars are standard deviation. True detection is the detection of the Projection-TAG expressed in a given sample transfected with one Projection-TAG plasmid, false detection is the detection of Projection-TAGs not expressed in the same samples, and the background is the detection of individual Projection-TAGs in the HEK sample with no transfection. **d**, Representative FISH images showing the detection of individual Projection-TAGs (BCs 1-12) in individual HEK samples each expressing a unique TAG or the HEK sample with no transfection. The bar plots show the average signal intensity from each Projection-TAG channel across a field of view with 1 mm by 1 mm. The average intensity is normalized by the max values of each sample and each channel. In HEK samples transfected with Projection-TAG plasmids, the dotted line and the associated number represent the false detection rate (average intensity of Projection-TAGs not expressed in a given sample), whereas the number on the right indicates the true detection rate (intensity from the Projection-TAG expressed in a given sample). In the HEK sample with no transfection, the dotted line and the associated number represent the background (average intensity of all Projection-TAGs). Scale bar: 20 µm. **e**, Bar plot showing average detection rate of Projection-TAGs across all samples, error bars are standard deviation. **f**, Representative images showing that photobleaching process removed the native Sun1-GFP fluorescence from HEK cells. Scale bar: 50 µm. **g**, Representative images showing that DNase incubation removed the fluorescent signal of Alexa-488 conjugated with *GAPDH* FISH probe in HEK cells. Scale bar: 50 µm. **h**, Representative images showing that photobleaching process significantly reduced the native Sun1-GFP fluorescence from mouse brain sections. Scale bar: 100 µm. **i**, Representative images showing that DNase incubation removed the fluorescent signals of Alexa-488 conjugated with the FISH probe targeting *GFP* in mouse brain sections. Scale bar: 100 µm.

**Extended Data Fig. 3 – related to Fig. 1.**
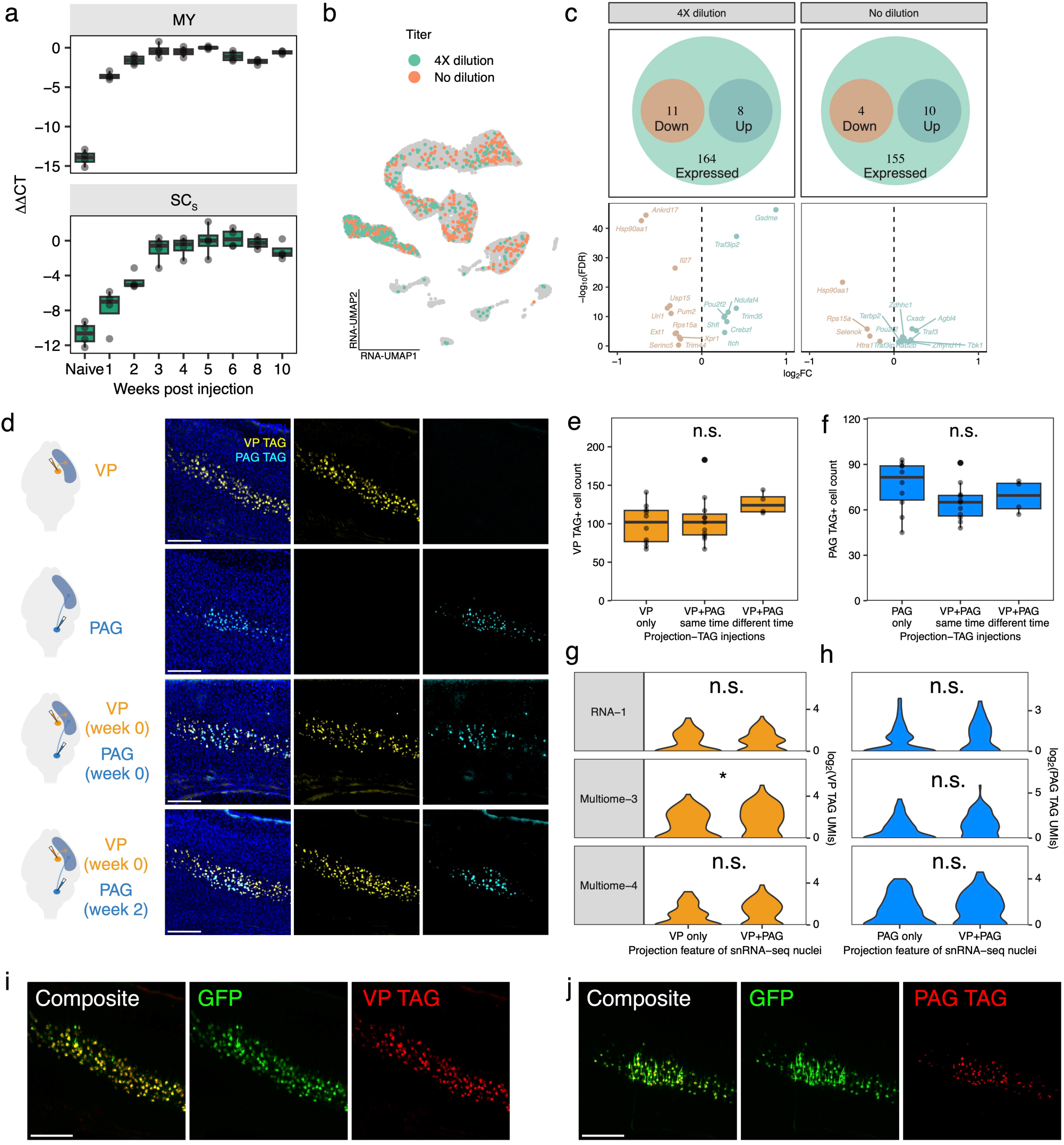
**a**, Box plot showing the time kinetics of Projection-TAG expression in the cortex. Animals received MY or SC_S_ injection with the Projection-TAG AAV (expressing TAG 2), cortical samples were collected at different time points post injection, and the Projection-TAG expression is measured using qPCR (n=4 for each time point and each injection). ΔCT is calculated as the CT value of TAG 2 minus that of GAPDH for each sample, ΔΔCT is calculated as the ΔCT at each time point minus the ΔCT at 4 weeks post-injection. **b**, RNA-UMAP of 10,000 nuclei (1,000 downsampled Projection-TAG+ nuclei from each titer group, colored by titer, and 8,000 downsampled BC-nuclei, colored grey). **c**, DE genes in response to viral infection. DE analysis was performed comparing the expression of genes annotated in the Gene Ontology list GO:0009615 (response to virus) between Projection-TAG+ nuclei and Projection-TAG-nuclei (downsampled to match the Celltype distribution and nuclei counts to the Projection-TAG+ nuclei) in each tier group. Top panels show the number of upregulated (avg_log2FC > 0) and downregulated (avg_log2FC < 0) viral responding genes among the viral responding genes expressed (average expression > 0.5) in snRNA-seq nuclei from each titer group. Fisher’s exact test was performed to test if viral responding genes are significantly enriched among DE genes (p = 0.81 and 0.77 for undiluted and diluted group, respectively). Bottom panels show the individual DE viral responding genes. **d**, Representative FISH images of medial part of the posterior parietal association area of the cortex (PTLp) visualizing Projection-TAGs injected into VP and PAG in a mouse receiving only VP injection (first row), a mouse receiving only PAG injection (second row), a mouse receiving VP and PAG injections at the same time (third row), and a mouse receiving PAG injection two weeks after VP injection (bottom row). Scale bar: 100 µm. **e**, Quantification of VP-TAG+ cells in medial PTLp from different injection groups (nine slices from three mice each group). *p* = 0.26, F(2,25) = 1.72, one-way ANOVA. **f**, Quantification of PAG-TAG+ cells in medial PTLp from different injection groups (9 slices from three mice each group). *p* = 0.27, F(2,25) = 1.4, one-way ANOVA. **g**, Violin plot showing the VP-TAG UMIs from snRNA-seq nuclei expressing only VP-TAG and co-expressing VP/PAG-TAGs in three snRNA-seq libraries that contain at least 60 nuclei co-expressing VP/PAG-TAGs. *p* = 0.99 (RNA-1), 0.01 (Multiome-3), and 0.06 (Multiome-3), student t-tests with Bonferroni correction. **h**, Violin plot showing the PAG-TAG UMIs from snRNA-seq nuclei expressing only PAG-TAG and co-expressing VP/PAG-TAGs in the snRNA-seq libraries shown in g. *p* = 0.5 (RNA-1), 0.1 (Multiome-3), and 0.6 (Multiome-3), student t-tests with Bonferroni correction. **i** and **j**, representative images showing the co-localization of GFP fluorescent signals with VP-TAG signal (i) or PAG-TAG signal (j) from FISH. Scale bar: 100 µm.

**Extended Data Fig. 4 – related to Fig. 1.**
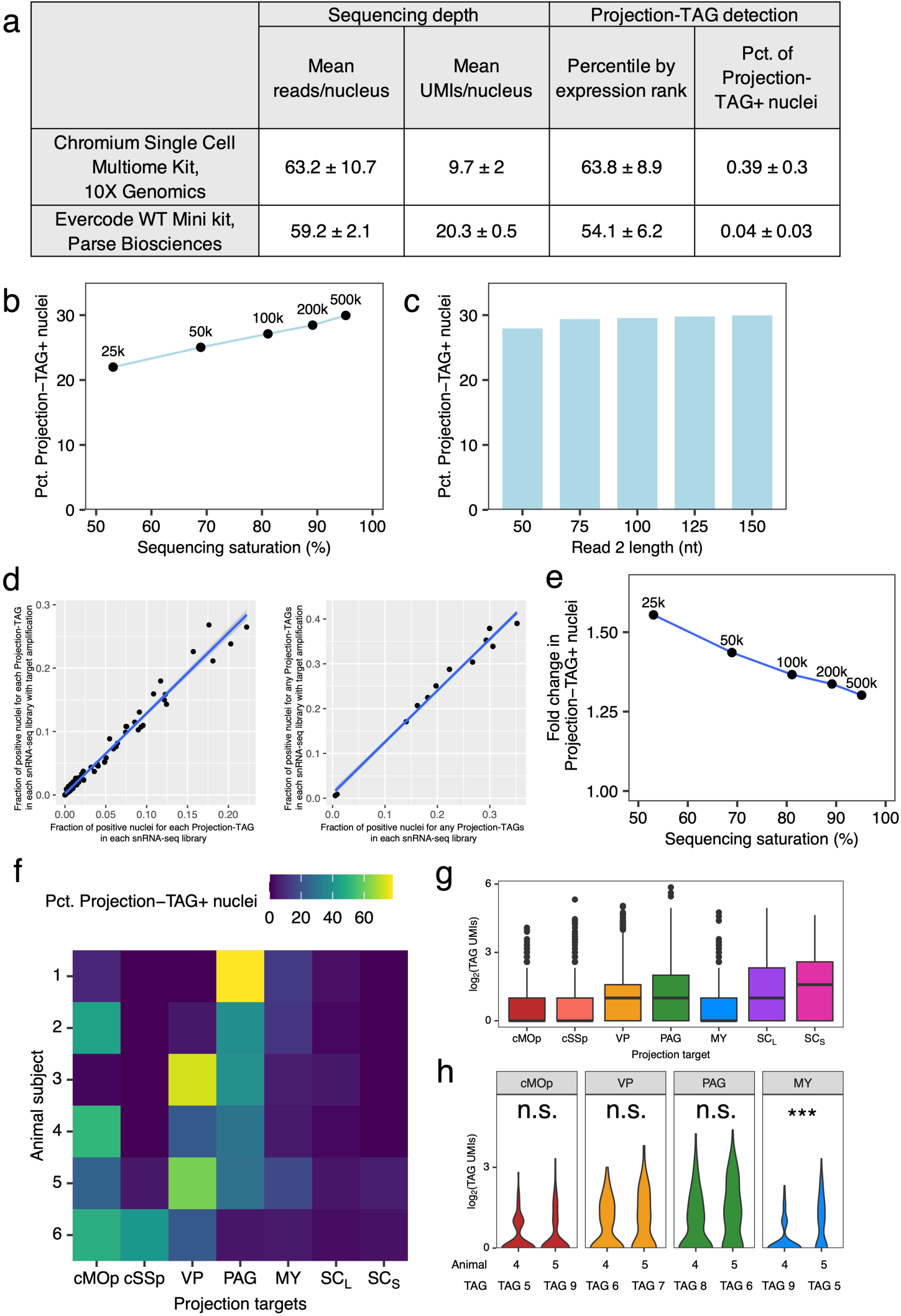
**a**, Key sequencing and Projection-TAG detection metrics from snRNA-seq libraries prepared using different commercial kits from nuclei of the same biological samples (n = 2 for each kit). **b**, Dot plot showing the effect of sequencing depth/saturation on the percentage of Projection-TAG+ nuclei in the snRNA-seq library that reached 500k reads/nucleus. Sequencing reads were randomly downsampled to match the sequencing reads/nucleus metric shown on the plot. **c**, Bar plot showing the effect of sequencing Read 2 length on the percentage of Projection-TAG+ nuclei in the snRNA-seq library shown in b. Sequencing reads were trimmed using “r2-length” parameter in cellranger. **d**, Detection of each Projection-TAG (left) and any Projection-TAGs (right) in snRNA-seq alone (x-axis) and in snRNA-seq library with target amplification (y-axis) in individual sequencing libraries. Blue line on each plot indicates the best fitted line. **e**, Dot plot showing the target amplification performance as fold change of Projection-TAG+ nuclei between snRNA-seq library with target amplification and snRNA-seq alone in the snRNA-seq library shown in b. Sequencing reads/nucleus metric is labeled on the plot **f**, Heatmap showing the percentage of Projection-TAG+ nuclei expressing each Projection-TAG (labeled as the injection site of the corresponding Projection-TAG on the row) from each animal. **g**, Box plot showing the Projection-TAG UMI count from snRNA-seq nuclei expressing each Projection-TAG (labeled as the injection site of the corresponding Projection-TAG on the row). **h**, Violin plots showing the Projection-TAG UMI count from snRNA-seq nuclei expressing the corresponding Projection-TAG from two animals (x-axis), in which different Projection-TAGs are injected into the same brain region for the regions shown on the plot. *p* = 0.08 cMOp, *p* = 0.12 VP, *p* = 0.65 PAG, \*\*\**p* = 1.5e-4 MY.

**Extended Data Fig. 5 – related to Fig. 1.**
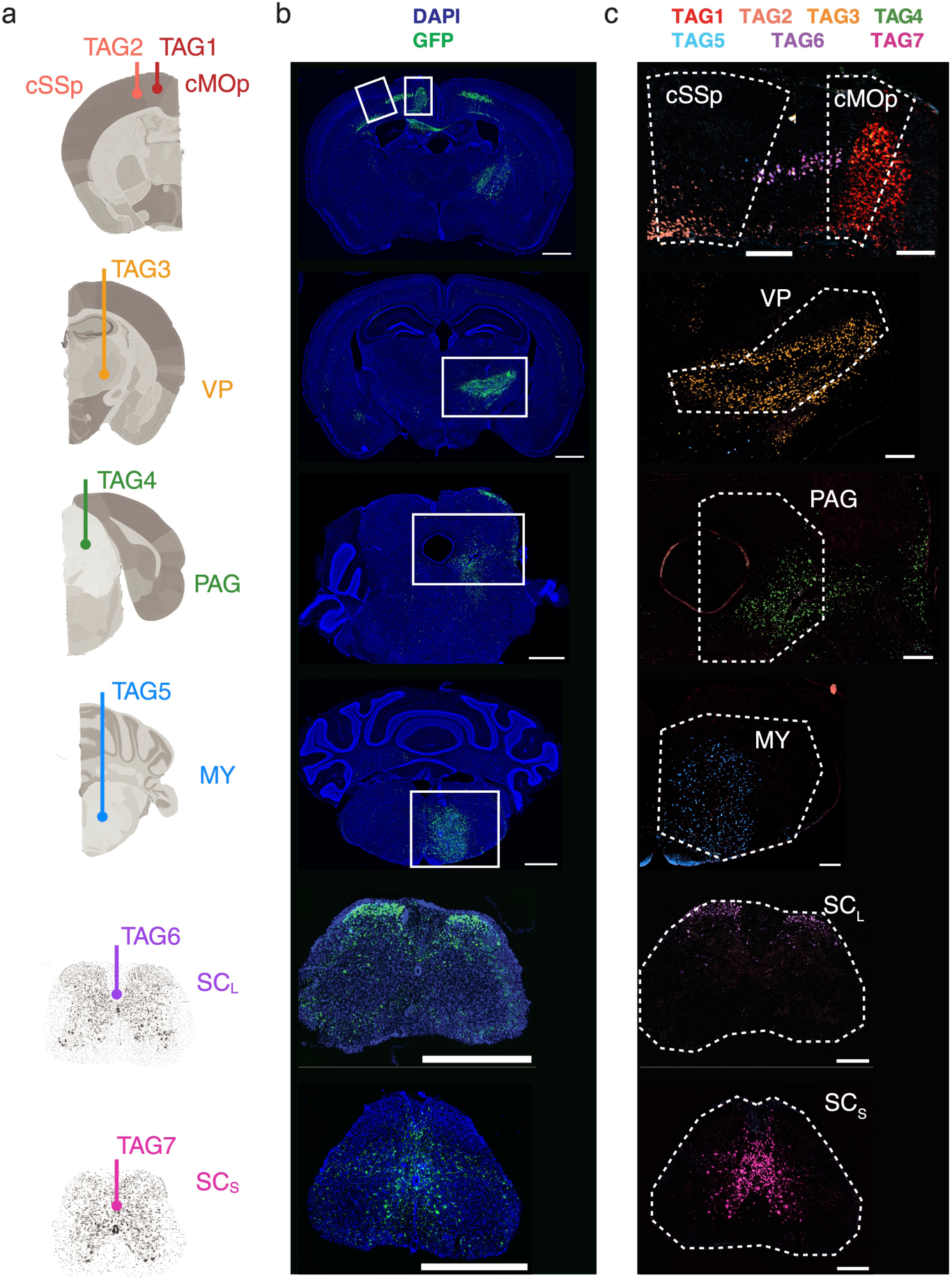
**a**, Schematics showing the stereotaxic injection of individual Projection-TAGs into each of the projection targets of the cortex. **b**, Representative images showing the native fluorescence of Sun1-GFP in the injection sites. Scale bar: 1 mm. **c**, Representative FISH images (zoom-in view of the region highlighted in b) showing the expression of Projection-TAGs in the injection sites. Multiplexed FISH was performed to visualize the Projection-TAGs (TAGs 1-7) in the projection targets of the cortex (highlighted with dotted lines). Scale bar: 300 µm.

**Extended Data Fig. 6 – related to Fig. 1.**
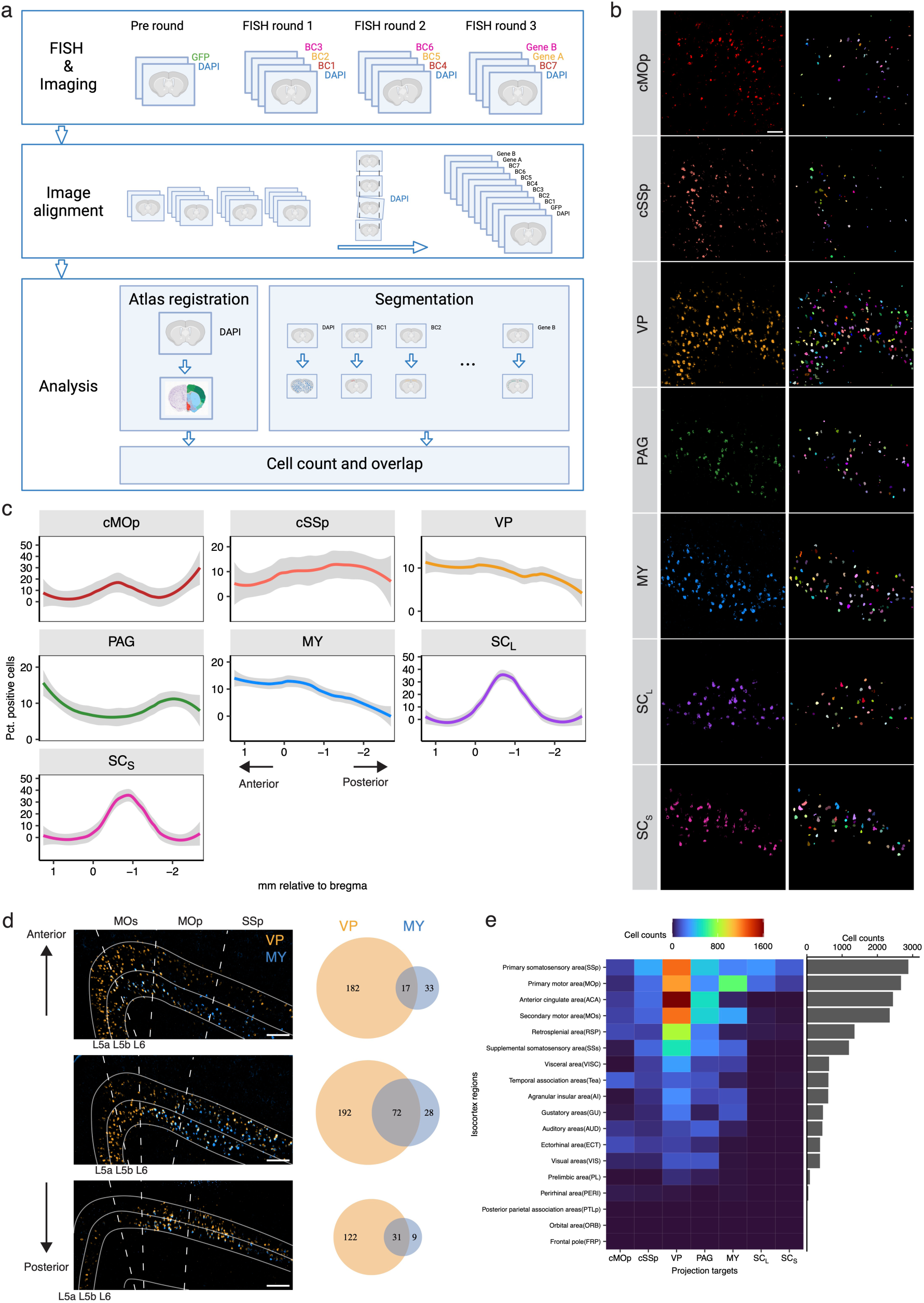
**a**, Workflow of multiplexed FISH experiment and imaging analysis. **b**, Left: representative FISH images showing the Projection-TAGs labeled projection neurons to each target (row) in the MOp. Right: cell segmentation of the projection neurons from the images on the left panels using machine-learning-based algorithms. Colors denote individual objects identified by the algorithms. **c**, Rostro-caudal distribution of neurons projecting to each target in ipsilateral cortex areas. Smooth lines were fitted to the quantification of 22 brain sections from two mice, and the grey ribbons show the standard error of the mean (SEM). **d**, Representative images of VP- and MY-projecting neurons labeled with Projection-TAGs in the select cortex areas from three brain sections (top: 1.26 mm, middle: 0.23 mm, bottom -0.95 mm relative to bregma). Venn diagrams show overlapping of cells projecting to both targets (*p* = 7.57e-14, 1.23e-44, 3.30e-41 for top, middle, and bottom diagrams, respectively. Hypergeometric tests). Scale bar: 200 µm. **e**, Heatmap showing the cell counts of neurons projecting to each target in each select cortex area.

**Extended Data Fig. 7 – related to Fig. 2.**
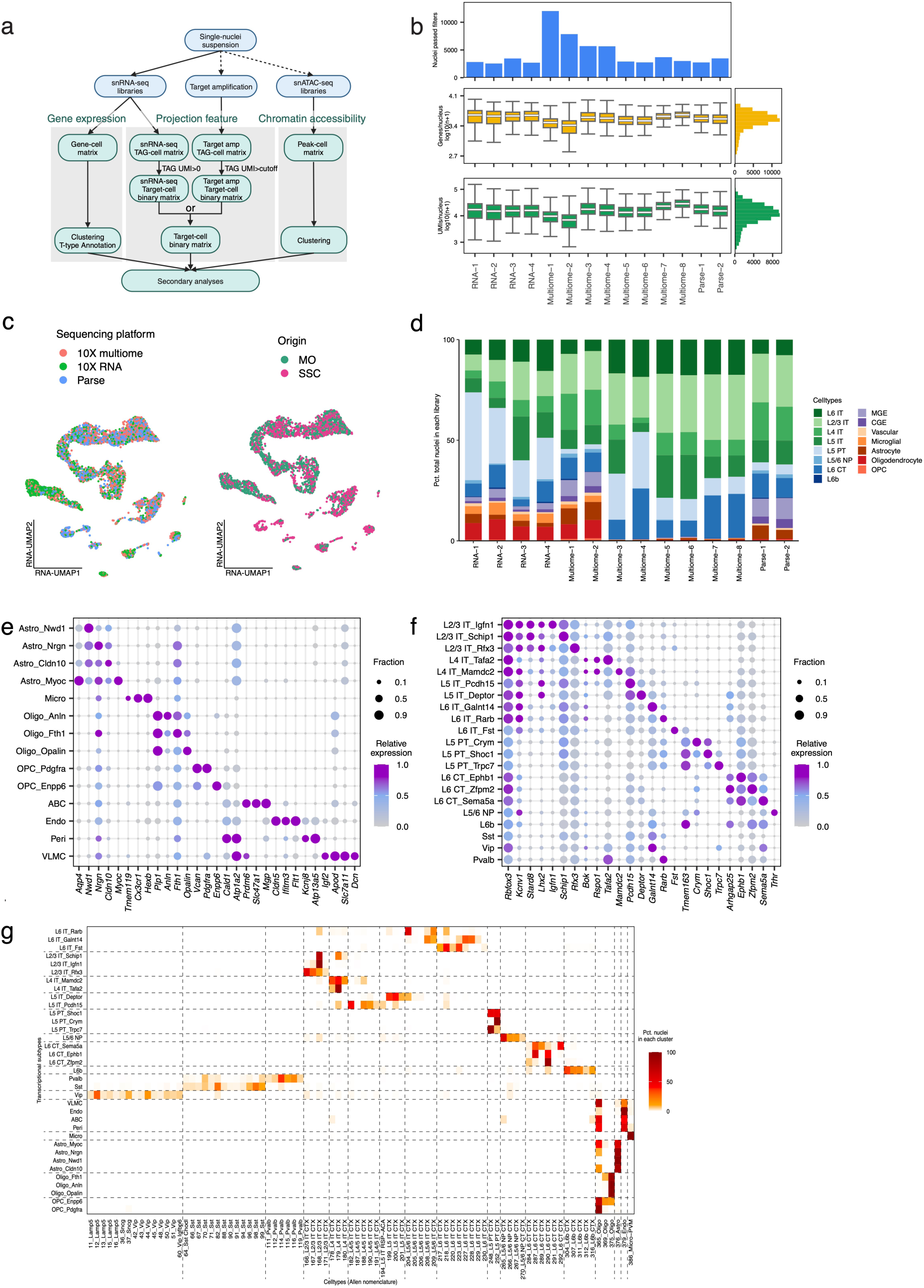
**a**, Workflow of the data analysis of the snRNA-seq and snATAC-seq and celltype and projection annotation. **b**, snRNA-seq library metrics. Top row displays number of nuclei passed quality control. Middle row displays box plots of number of genes per nucleus (log10 transformed) and the bottom row displays the number of UMIs per nucleus. Boxes indicate quartiles and whiskers are 1.5-times the interquartile range (Q1-Q3). The median is a white line inside each box. The distribution is aggregated across all samples and displayed on the horizontal histogram. **c**, RNA-UMAP of 1,000 downsampled nuclei per group. Nuclei were colored by sequencing kits (left) or the origin of brain regions (right). **d**, Composition of transcriptional celltypes in each snRNA-seq library. **e** and **f**, Dot plots showing the marker genes used to annotate subtypes of non-neuronal cells (e) and neuronal cells (f). **g**, Heatmap showing the percentage of snRNA-seq nuclei in each cluster (row) assigned by anchoring analysis to the celltypes (column) of a published scRNA-seq dataset of mouse MOp and SSp^123^.

**Extended Data Fig. 8 – related to Fig. 2.**
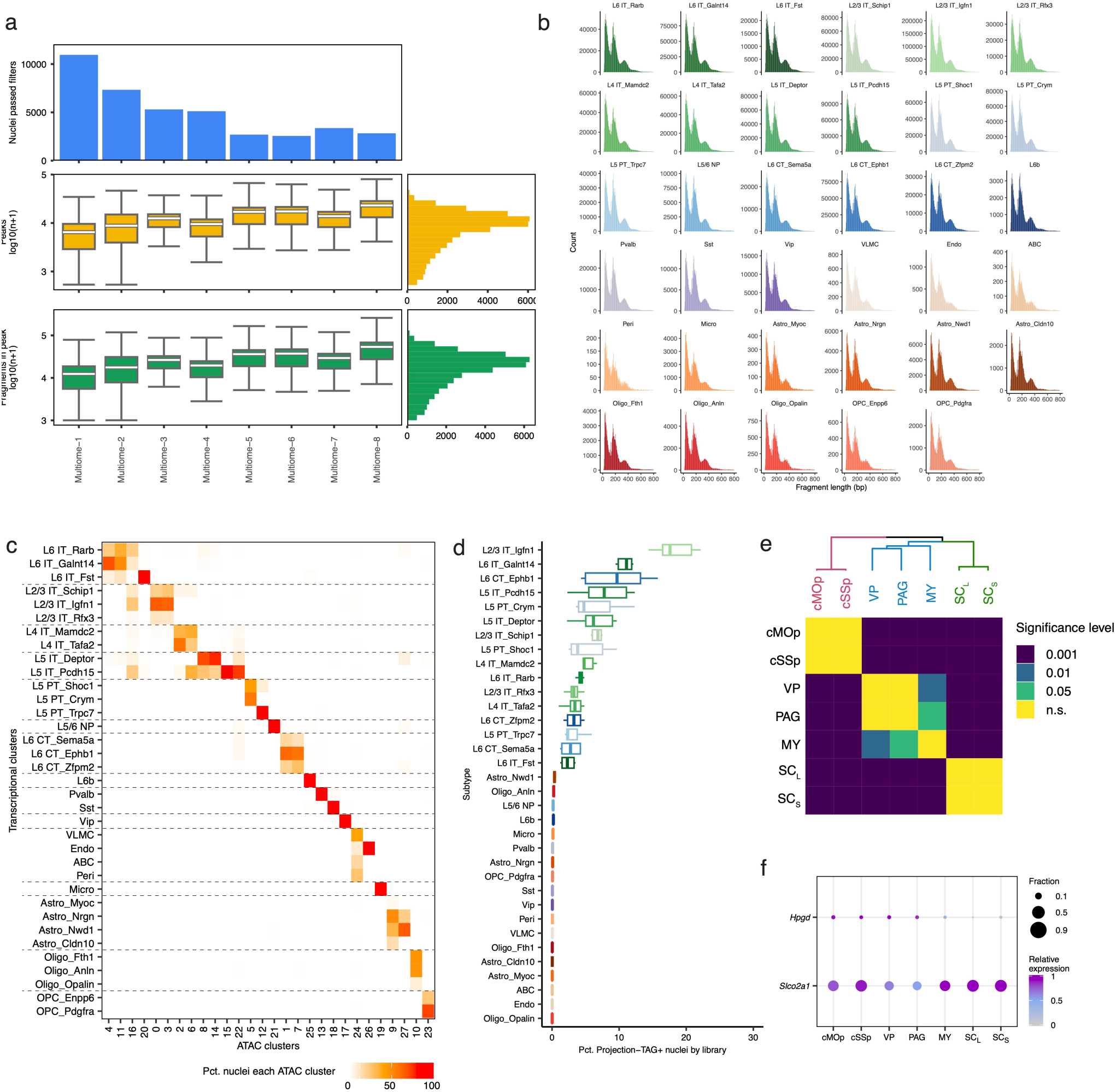
**a**, snATAC-seq library metrics. Top row displays number of nuclei passed quality control. Middle row displays box plots of number of peaks per nucleus (log10 transformed) and the bottom row displays the number of fragments that overlap with peaks per nucleus. Boxes indicate quartiles and whiskers are 1.5-times the interquartile range (Q1-Q3). The median is a white line inside each box. The distribution is aggregated across all samples and displayed on the horizontal histogram. **b**, Distribution of snATAC-seq fragment lengths in nuclei grouped by transcriptional subtypes. **c**, Correspondence of ATAC clusters (row) and transcriptional subtypes (column) of nuclei profiled combinatorially by snRNA-seq and snATAC-seq. Plot displays the percentage of nuclei within each ATAC cluster that is assigned to each transcriptional subtype. **d**, Box plot showing the percentage of Projection-TAG+ nuclei in each snRNA-seq library in each subtype. **e**, Heatmap showing the significance levels of the Bonferroni-corrected *p* values of pairwise hypergeometric tests of subtype distribution, shown in Fig. 2g, between any of the two projections. Hierarchical clustering was performed based on the subtype distribution for each projection. **f**, Dot plot showing the expression of *Hpgd* and *Slco2a1* in individual projections.

**Extended Data Fig. 9 – related to Fig. 3.**
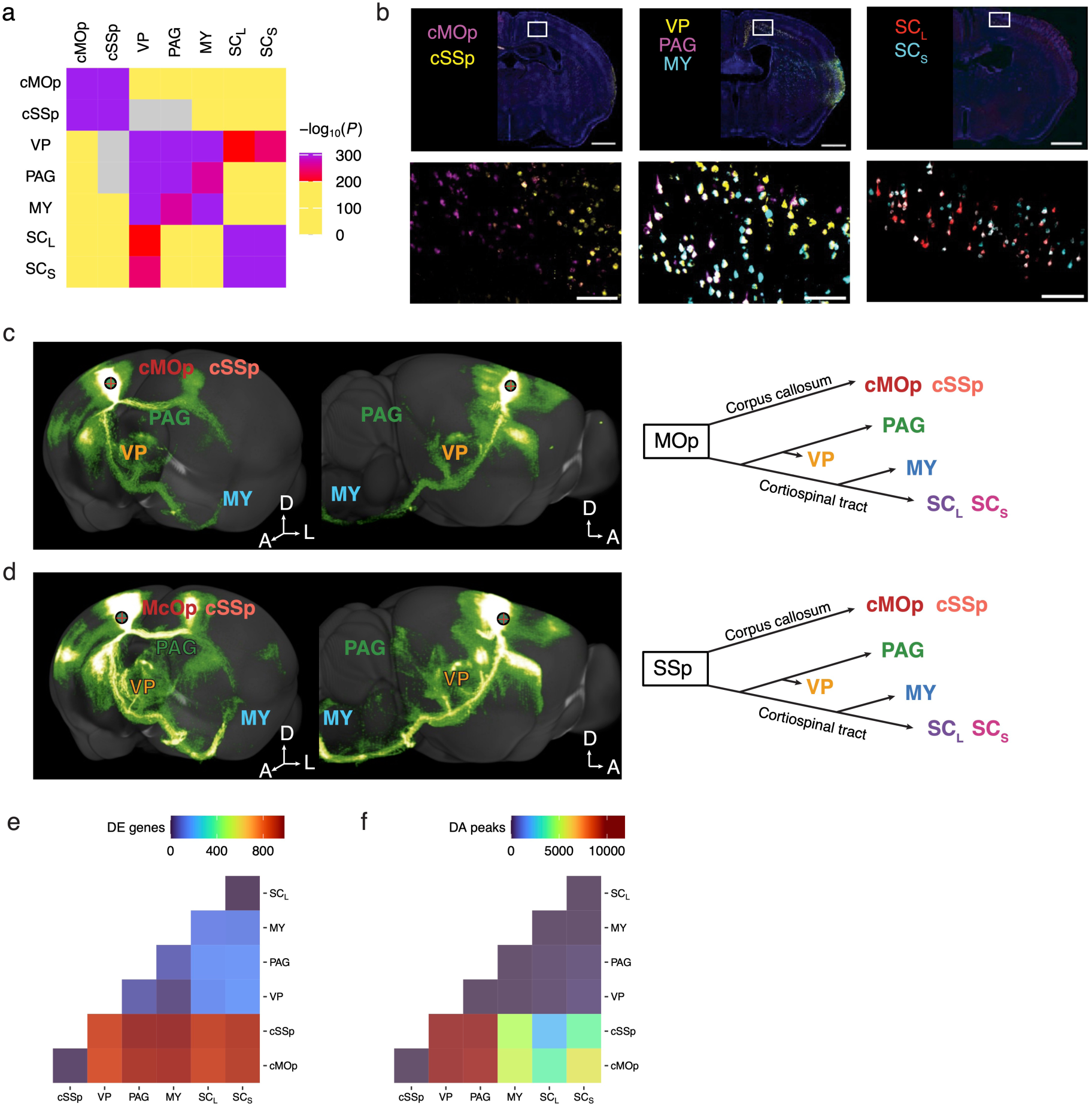
**a**, Heatmap showing the Bonferroni-corrected *p* values from pairwise hypergeometric tests of overlapping between any of the two projections. Two projections were considered highly significantly overlapping with *p* < 1e-200 (shown as red to purple on the heatmap). **b**, Representative FISH images showing projection neurons labeled by Projection-TAGs in the coronal section showing motor cortex from different anterior-posterior sections (top) and zoomed-in images (bottom) of the highlighted region demonstrate expression of two unique Projection-TAGs injected into downstream cMOp and cSSp; middle: demonstrate expression of three unique Projection-TAGs injected into the VP, PAG, and MY; right: demonstrate expression of two unique Projection-TAGs injected into SC_L_ and SC_S_). Scale bar: 1 mm (top), 100 µm (bottom). **c**, Tracing the brain-wide axon collaterals of MOp neurons. Screenshots acquired from Allen Brain Atlas data portal (https://connectivity.brain-map.org/projection/experiment/100141563)^125^ were displayed on the left and a summary diagram of the anatomical hierarchy of axonal projections from MOp to the seven target regions was shown on the right. **d**, Tracing the brain-wide axon collaterals of SSp neurons. Images acquired from Allen Brain Atlas data portal (https://connectivity.brain-map.org/projection/experiment/112935169)^125^. were displayed on the left and a summary diagram of the anatomical hierarchy of axonal projections from SSp to the seven target regions was shown on the right. **e** and **f**, Heatmap showing the number of DE genes (FDR<0.05, e) and DA peaks (FDR<0.05, f) between any two projections.

**Extended Data Fig. 10 – related to Fig. 3.**
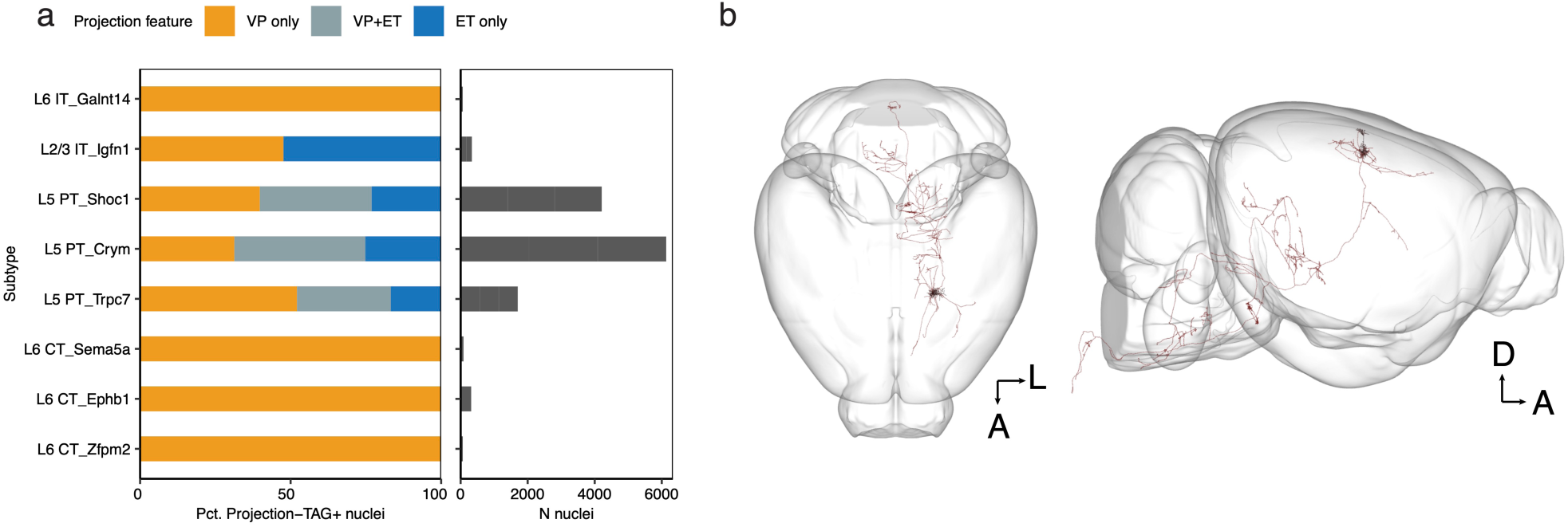
**a**, Distribution of projection feature to VP and/or other ET targets (defined in Fig. 3h-j) in Projection-TAG+ nuclei in each subtype.. **b**, A single neuron (MouseLight AA0923) whose cell body reside in the MOp L5 and axonal end points terminating in thalamus, PAG, MY, an axon follow through the corticospinal tract. Screenshots were acquired from the Janelia MouseLight data portal (http://ml-neuronbrowser.janelia.org)^69^.

**Extended Data Fig. 11 – related to Fig. 4.**
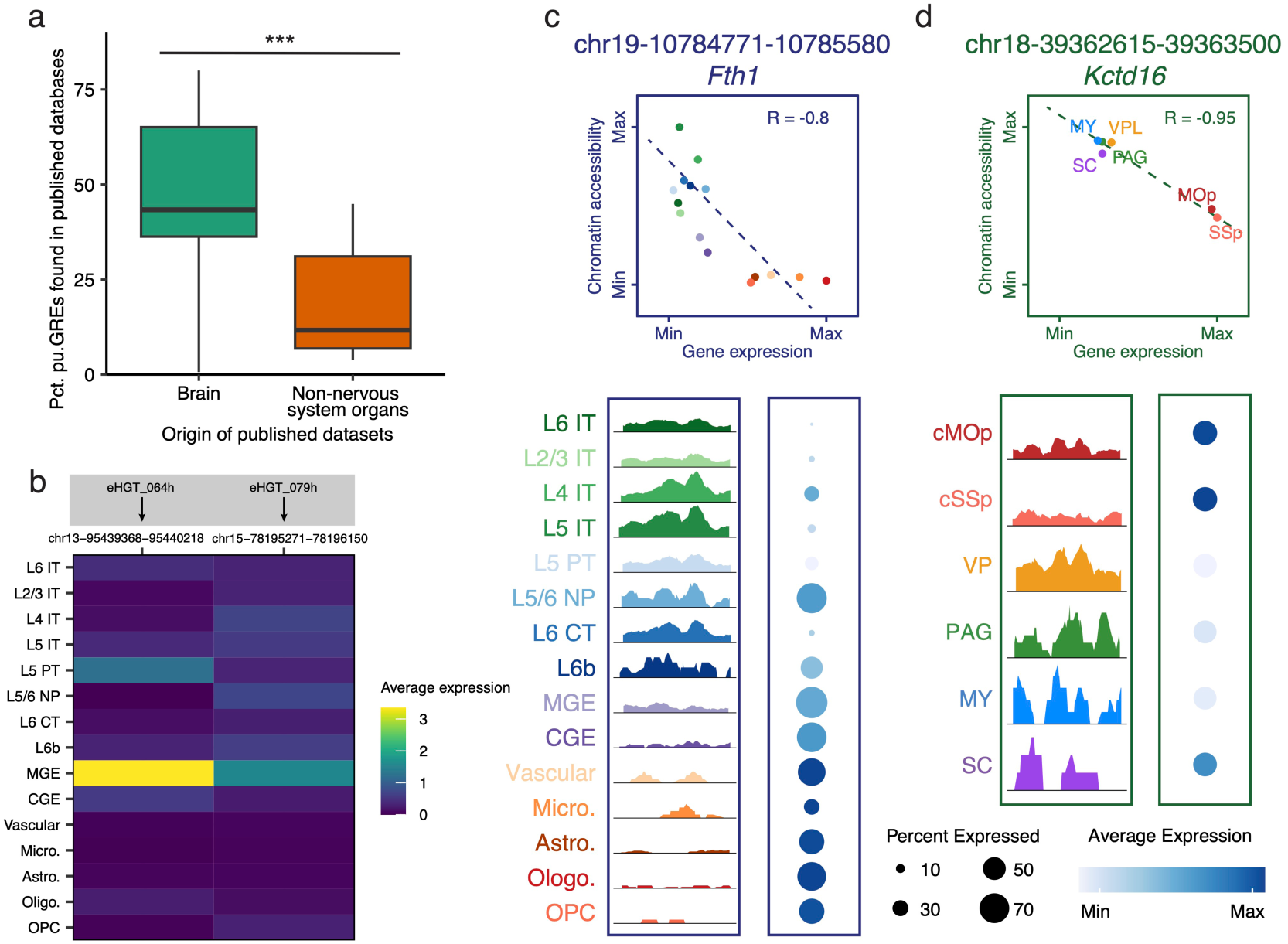
**a**, Box plot showing the overlapping of putative GREs identified from this study to the putative GREs identified by studies from the ENCODE consortium^83,126^. Overlap is significantly higher with studies of the brain compared to those of non-brain organs (\*\*\**p* = 8.26e-05, t-test). Brain studies: ENCSR422CGZ, ENCSR694ZBB, ENCFF426KUB, and ENCSR410XMU. Studies of organs that are not the nervous system: ENCSR052AZT (stomach), ENCSR100TMZ (lung), ENCSR296UQL (liver), and ENCSR180OIF (limb). **b**, Heatmap showing the chromatin accessibility of two celltype-specific pu.Enhancers in individual Transcriptional subtypes. The peaks of the two pu.Enhancers overlap with the genomic coordinates liftover from two functional enhancers previously identified based on human snATAC-seq data^77^. Labels in the grey boxes show the functional enhancers and the cell types in which the enhancers exhibited highest activity in experimental validation from the previous study. **c**, Top: Scatter plot showing the correlation between the accessibility of peak chr19-10,784,771-10,785,580 and the expression of gene *Fth1* in individual transcriptional subtypes (chromatin accessibility and gene expression are normalized to their max values). The dotted line represents the line of best fit. bottom left: chromatin accessibility at the genomic locus of chr19-10,784,771-10,785,580, displayed as the average fraction of transposase-sensitive fragments per nucleus in each subtype (grouped by 50 bins per displayed genomic region). Accessibility at each locus (y axis) is scaled to the max value across all cell types. Bottom right: expression of *Fth1* in each celltype. Colors indicate Transcriptional subtypes. **d**, Top: Scatter plot showing the correlation between the accessibility of peak chr18-39,363,615-39,363,500 and the expression of the gene *Kctd16* in nuclei positive for individual TAGs (chromatin accessibility and gene expression are normalized to their max values). bottom left: chromatin accessibility at the genomic locus of chr18-39,363,615-39,363,500, displayed as the average fraction of transposase-sensitive fragments per nucleus in each projection (grouped by 50 bins per displayed genomic region). Bottom right: expression of *Kctd16* in each projection. Colors indicate nuclei positive for individual TAGs. Nuclei positive for SC_L_- and SC_S_-TAGs are combined for visualization due to low cell number.

**Extended Data Fig. 12 – related to Fig. 5.**
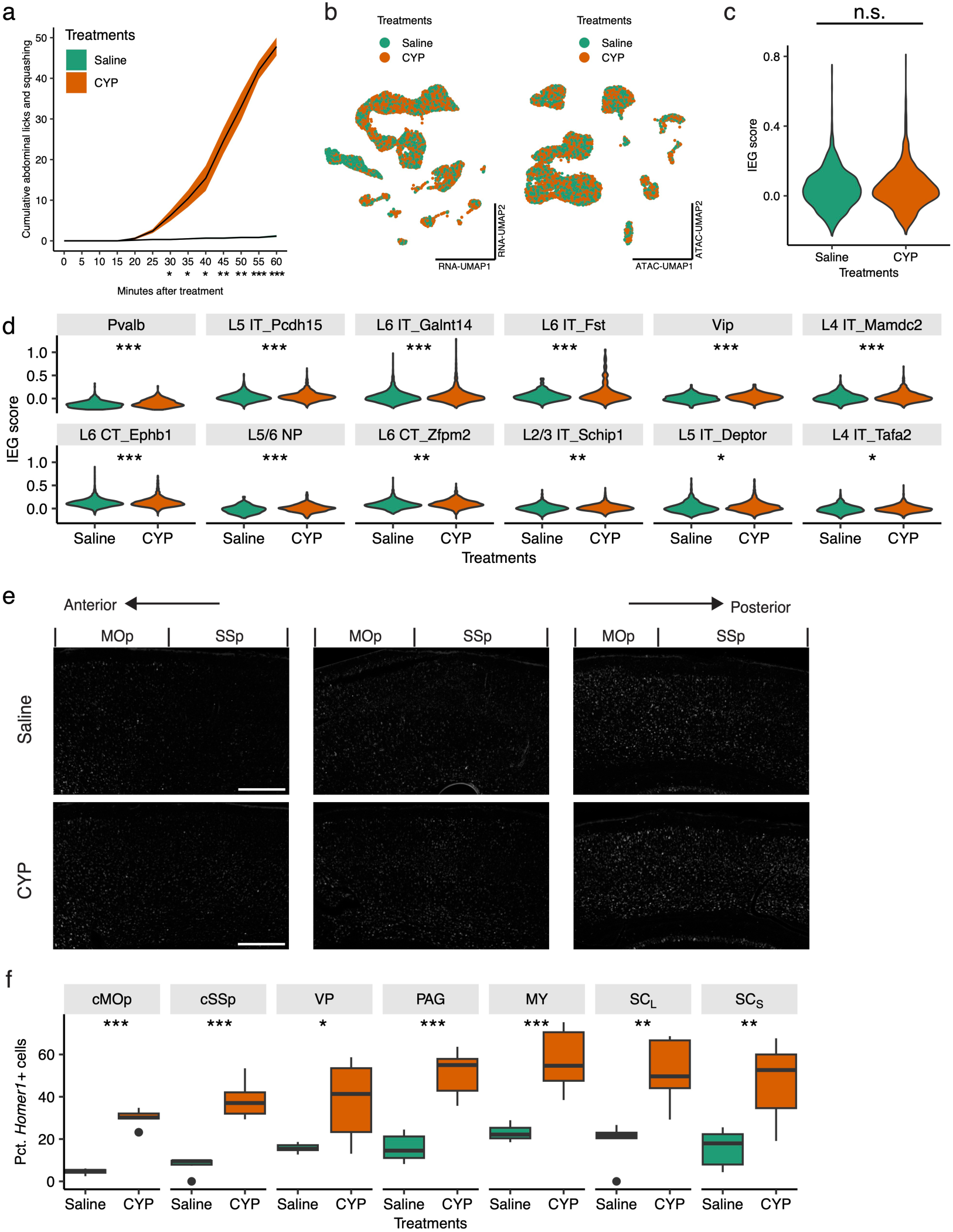
**a**, Cumulative scores of quantified behaviors between mice treated with saline and CYP. CYP treatment resulted in significant increases in the spontaneous behaviors like abdominal licking and squashing behaviors compared to saline treatment. Ribbons show SEM. ** *p* = 0.0034, one-way ANOVA. p = NaN (0 min), NaN (5 min), NaN (10 min), NaN (15 min), 0.25 (20 min), 0.07 (25 min), *0.05 (30 min), *0.04 (35 min), *0.04 (40 min), **0.001 (45 min), **0.006 (50 min), ***0.0005 (55 min), ***0.0003 (60 min), respectively, post-hoc one-sided t-test, n=5-6 mice/group. **b**, RNA-UMAP (left) and ATAC-UMAP (right) of 5,000 downsampled nuclei from each treatment group, colored by treatments. **c**, Violin plot showing IEG scores of all snRNA-seq nuclei from CYP and saline. *p* = 0.16. **d**, Violin plots showing IEG score of nuclei between treatments from CYP-activated transcriptional subtypes. \*\*\**p* = 2.6e-7 Pvalb, \*\*\**p* = 6.2e-7 L5 IT_Pcdh15, \*\*\**p* = 1.3e-6 L6 IT_Galnt14, \*\*\**p* = 1.5e-6 L6 IT_Fst, \*\*\**p* = 4.6e-5 Vip, \*\*\**p* = 1.5e-4 L4 IT_Mamdc2, \*\*\**p* = 3.8e-4 L6 CT_Ephb1, \*\*\**p* =3.8e-4 L5/6 NP, \*\**p* = 1.3e-3 L6 CT_Zfpm2, \*\**p* = 4.6e-3 L2/3 IT_Schip1, \**p* = 0.02 L5 IT_Deptor, \**p* = 0.04 L4 IT_Tafa2. **e**, Representative FISH images showing *Fos* expression in select cortex areas (top: 1.33 mm, middle: -0.95 mm, bottom -1.67 mm relative to bregma). Scale bar: 400 µm. **f**, Bar plot showing the percentage of Homer1+ cells in each projection group from FISH analysis. Nine slices from three mice per group, \*\*\**p* = 6.7e-8 cMOp, \*\*\**p* = 2.9e-6 cSSp, \**p* = 0.02 VP, \*\*\**p* = 1.2e-5 PAG, \*\*\**p* = 1.9e-4 MY, \*\**p* = 0.002 SC_L_, \*\**p* = 0.006 SC_S_.

## Notes

### Competing Interest Statement

The authors have declared no competing interest.

### Summary of Updates

We have update figure 3 analysis and updated the manuscript discussion.

